# Whole genome sequence accuracy is improved by replication in a population of mutagenized sorghum

**DOI:** 10.1101/095513

**Authors:** Charles Addo-Quaye, Mitch Tuinstra, Nicola Carraro, Clifford Weil, Brian P. Dilkes

## Abstract

The accurate detection of induced mutations is critical for both forward and reverse genetics studies. Experimental chemical mutagenesis induces relatively few single base changes per individual. In a complex eukaryotic genome, false positive detection of mutations can occur at or above this mutagenesis rate. We demonstrate here, using a population of ethyl methanesulfonate (EMS) treated *Sorghum bicolor* BTx623 individuals, that using replication to detect false positive induced variants in next-generation sequencing data permits higher throughput variant detection with greater accuracy. We used a lower sequence coverage depth (average of 7X) from 586 independently mutagenized individuals and detected 5,399,493 homozygous SNPs. Of these, 76% originated from only 57,872 genomic positions prone to false positive variant calling. These positions are characterized by high copy number paralogs where the error-prone SNP positions are at copies containing a variant at the SNP position. The ability of short stretches of homology to generate these error prone positions suggests that incompletely assembled or poorly mapped repeated sequences are one driver of these error prone positions. Removal of these false positives left 1,275,872 homozygous and 477,531 heterozygous EMS-induced SNPs which, congruent with the mutagenic mechanism of EMS, were greater than 98% G:C to A:T transitions. Through this analysis we generated a database of sequence indexed mutants of Sorghum. This collection contains 4,035 high impact homozygous mutations in 3,637 genes and 56,514 homozygous missense mutations in 23,227 genes. Each line contains, on average, 2,177 annotated homozygous SNPs per genome, including seven likely gene knockouts and 96 missense mutations. The number of mutations in a transcript was linearly correlated with the transcript length and also the G+C count, but not with the GC/AT ratio. Analysis of the detected mutagenized positions identified CG-rich patches, and flanking sequences strongly influenced EMS-induced mutation rates. Our method for detecting false-positive induced mutations is generally applicable to any organism, is independent of the choice of *in silico* variant-calling algorithm, and is most valuable when the true mutation rate is likely to be low, such as in laboratory induced mutations or somatic mutation detection in medicine.

## INTRODUCTION

Forward and reverse genetics are ubiquitous methods for gene function discovery (Alonso & Ecker, 2006; Nordborg & Weigel, 2008; Page & Grossniklaus, 2002; Schneeberger & Weigel, 2011). The principal step in these methods is the induction of mutations in the germline of an organism using various categories of high energy (such as UV light, X-rays, gamma rays and fast neutrons) or chemical mutagens (such as EMS, MMS and ENU). The objective is the production of genetic diversity in a population and identification of phenotypic variants (Waugh, Leader, McCallum, & Caldwell, 2006). The chemical ethyl methanesulfonate (EMS) is extensively used for mutagenesis (Sega, 1984), and predominantly induces G:C to A:T transitions in the genome of a treated organism (Coulondre & Miller, 1977; Krieg, 1963; Loveless, 1958; Loveless, 1969; Prakash & Sherman, 1973; Singer, Fraenkel-Conrat, & Kusmierek, 1978; Westergaard, 1957). The accurate detection of these induced mutations is critical to establish evidence of gene function.

The ascendancy of cost-effective DNA sequencing methods has enhanced our capacity to detect induced polymorphisms (Metzker, 2010) and several sequencing-based methods have been proposed for detecting induced polymorphisms in plant genetic studies (Nordborg & Weigel, 2008; Nordström et al., 2013; Rowan, Weigel, & Koenig, 2011; Schlötterer, Tobler, Kofler, & Nolte, 2014; Schneeberger & Weigel, 2011; Schneeberger, 2014). Despite successes and widespread use of these methods, their accuracy is impacted negatively by the inherently higher error rates of next-generation sequencing (NGS) technologies (Nakamura et al., 2011; Nielsen, Paul, Albrechtsen, & Song, 2011), disparities in variant-calling methods (Cheng, Teo, & Ong, 2014; O’Rawe et al., 2013), qualities of *in silico* reconstructed reference genome sequences (Cheung et al., 2003), and the genome complexity of target organisms (Estivill et al., 2002; Fredman et al., 2004; Robasky, Lewis, & Church, 2014). Improving the accuracy of methods for detecting induced polymorphism in a cost-effective manner would be valuable for gene function discovery studies outside of genetic model species, which are often characterized by small genome sizes.

Sorghum is a fully-sequenced panicoid grass and an important source of food, feed and potential biomass for biofuel production (Paterson, 2008). It is well-adapted to hot, semi-arid conditions, nutrient-poor soils, and has particularly high drought tolerance relative to other cultivated cereals (Paterson et al., 2009). At 0.8 Gb the sorghum genome is more compact than maize (2.3Gb) and sugarcane (10Gb) and has not undergone a recent polyploidy event (Arumuganathan & Earle, 1991; Peterson et al., 2002; Price et al., 2005; Swigoňová et al., 2004). It thus offers a potentially simpler system in which to discover mechanisms of drought tolerance, plant nutrition, weed biology, evolution of C4 photosynthesis, and biofuel research. Recent forward genetics studies using EMS mutagenesis have provided key insights into the function of genes involved in insect-herbivory defense, cell wall and lignin biosynthesis, protein digestibility, leaf vein density, and plant height (Blomstedt et al., 2012; Koegel et al., 2013; Krothapalli et al., 2013; Pedersen, Bean, Graybosch, Park, & Tilley, 2005; Peters, Jenks, Rich, Axtell, & Ejeta, 2009; Petti et al., 2013; Petti, Hirano, Stork, & DeBolt, 2015; Rizal et al., 2015; Sattler et al., 2014; Scully et al., 2015; Wu, Yuan, Guo, Holding, & Messing, 2013; Xin, Wang, Burow, & Burke, 2009). The recent characterization of mutations in pooled samples derived from a population of 256 EMS-mutagenized sorghum BTx623 lines is a major step towards improving resources for sorghum genetics (Jiao et al., 2016). The availability of such large-scale collections of high quality and well-annotated mutations will play a pivotal role in addressing key questions in plant science research.

We have created an additional EMS-mutagenized sorghum population of ~12,000 lines and present here the large-scale detection and analysis of single nucleotide polymorphisms (SNPs) and insertions/deletions (INDELs) derived from low coverage, next-generation sequencing of 586 of these individuals. We treated these individuals as a population of replicates of sequencing error to remove false positive SNP calls. This cost-effective and accurate method effectively detects bona fide EMS-induced mutations and eliminates false positive calls. We characterized the predicted impact of these mutations on sorghum gene function and created a database of mutation descriptions, the corresponding altered genes and the gene function descriptions of their homologs in Arabidopsis and maize. Our method can be applied generally to genetic studies in most organisms, and is independent of the *in silico* variant-calling method utilized. We anticipate this valuable genetic resource and analytical approach will be useful to geneticists, plant breeders and the general plant science community.

## MATERIALS AND METHODS

### Mutagenesis and DNA preparation

Seed of the reference sorghum BTx623 genotype were treated with EMS (Sigma-Aldrich) by imbibing the seed overnight in distilled water at 20°C with shaking (150 rpm) and then soaking in 1L of 45mM EMS in 10mM KH_2_PO_4_ (pH 7.0) at 20°C for 6h shaking at 150 RPM. Seed were then washed ten times with 1L of distilled water and allowed to air dry at room temperature prior to planting. M_0_ seed were planted in an isolation block at approximately 20 seeds per meter. Unmutagenized BTx623 rows were interspersed with mutagenized seed to serve as pollen parents to improve seed set. Plants were allowed to open pollinate and one head was harvested from ~12,000 mutagenized plants, dried and threshed individually to produce M_1_ seed. A total of 20 M_1_ seeds were then planted for 7,304 lines. The tenth plant in each plot was self-fertilized to produce M_2_ seed. A total of 40 M_2_ seeds from 4,800 lines were planted to score for segregating phenotypes. M_2_ plants in each plot were self-fertilized to produce M_3_ seed. M_3_ seeds for each line were planted in the greenhouse. For each line, adult leaf tissue was harvested from a single plant for DNA preparation and stored at −20°C; the same plant was then self-fertilized to produce M_4_ seed. All DNA was prepared using a CTAB extraction as described by Xin and Chen (2012) except that incubations were done at 55°C rather than 60°C.

### Sequencing protocols

DNA was processed using the Illumina TruSeq DNA PCR-Free LT Library Preparation protocol, to make single indexed, 550bp insert libraries. These were titred individually using an Applied Biosystems Step One kit with Kapa Biosystems Illumina Library Quantification Kit reagents. After flowcell clustering using a TruSeq PE Cluster Kitv3-cBot-HS kit, sequence data was generated as paired 101 base reads by an Illumina HiSeq 2500 using SBS v3-HS reagents. “Stage 1” pools of 24 samples with compatible indices were created and a single lane of each pool was sequenced. Based on demultiplexing results from the initial lane of sequence, the library titres were refined and a new, “stage 2” pool created to correct for excesses/deficiencies in numbers of reads per library from the first lane. Two lanes of the stage 2 pool sequence were then run. Reads from all three runs were merged for each library.

### Next-generation sequencing reads mapping

Illumina HiSeq paired-end sequencing reads were mapped to version 2.1 of the sorghum genome reference sequences (Paterson et al., 2009) downloaded from the Phytozome (version 9.1) web portal (Goodstein et al., 2012). Reads alignment was performed using the BWA program for short reads alignment (Li & Durbin, 2009; version 0.6.2) and the variant calls were computed using the SAMtools software suite (Li et al., 2009; version 0.1.18). Sequenced reads and the reference genome sequences were indexed using the BWA *index* command with parameter “-a bwtsw”, and the SAMtools *faidx* command with default parameters, respectively. Suffix array co-ordinates for the NGS reads were generated using the BWA *aln* command (in multithreaded mode) with parameter “-t 8”. Alignments in the SAM format were generated using the BWA *sampe* command with parameter options: ‘ -P -r “@RG\tID:SAMPLEID\tSM:SAMPLENAME\tPL:Illumina”’. The SAM-formatted alignments were converted into binary BAM format using the SAMtools *view* command with parameter options: “-b -h -S -o”, and then sorted and indexed using the default parameters of the SAMTools *sort* and *index* commands, respectively. Genome mapping statistics were generated from the BAM-formatted files using the SAMtools *flagstat* command with default parameters. Genome coverage was estimated from the alignments in the BAM files using the combination of the SAMtools *view,* and the *genomeCoverageBed* command in the BEDTools genome arithmetic software suite (Quinlan & Hall, 2010; version 2.17.0). Command line parameters invoked for the SAMtools *view* and BEDTools *genomeCoverageBed* commands were “-u -q 0” and “-ibam stdin -g”, respectively. The output of the SAMtools *view* command was redirected to the BEDTools *genomeCoverageBed* command using a UNIX piping construct. Insert sizes for the paired-end reads were estimated using the *CollectInsertSizeMetrics* command in the Picard software package (http://broadinstitute.github.io/picard; version 1.80), with parameter option “VALIDATION_STRINGENCY=LENIENT”. Sample scripts to reproduce these analyses are provided in Supplementary Data 21.

### Detection of genomic positions with non-reference alignments

Whole genome variant calling started with the detection of positions with statistical support of non-reference sequence in each sequenced sorghum individual. We used the SAMtools *mpileup* command to compute mapping information for all genomic positions in every BAM file. We selected the following command line parameters to ensure that only DNA bases in sequenced reads with basecall qualities of at least 20 were used in variant detection: “-Q 20 -P Illumina -uf”. The *mpileup* output resulted in a Binary Call Format (BCF) file, which was redirected to the BCFtools *view* program. To produce a subset containing only the possible variant positions we used the BCFtools *view* command with parameter options: “-vcg” to output a variant call (VCF) file limited to likely variants only. This file contains bona fide SNP and indel positions, as well as false positive variant calls which satisfied these low stringency criteria that accept any position at which any read returned a non-reference alignment to be a variant position.

### SNPs filtering and refinement

The output of the variant calling stage was refined in multiple steps. We initially used the *varFilter* command (with non-default parameter option: “-D 100”) in the SAMtools *vcfutils.pl* Perl utility script to remove variants with low quality score (less than 10), low read coverage (read depth less than two), and possibly derived from repetitive origins (read depth greater than 100). We refer to this initially filtered SNP as the “Repeat-Filtered” SNPs. Next we used the UNIX *grep* utility (with parameter: -v “INDEL;”) to filter indel variant calls. We then used the *length* function in the UNIX *awk* utility to filter SNP variants in genomic positions with more than one non-reference SNP. We then used the SnpSift component (Cingolani, Patel, et al., 2012) in the SnpEff java package (Cingolani, Platts, et al., 2012; version 3.1) for further SNP variants filtering. Using the SnpSift *filter* command with query string “(QUAL>=20) & (isHom (GEN[0])) & (isVariant(GEN[0]))”, we retained homozygous SNPs with variant call quality of at least 20. This set of filtered SNPs we designate as the homozygous “Q20” SNPs. These SNPs were further filtered using the SnpSift *filter* command with query string “(((DP4[0]=0) & (DP4[1]=0)) & ((DP4[2]>0) & (DP4[3]>0)))” to retain SNPs with at least one non-reference read mapping to both the forward and reverse strands of the reference genome, that had no reference reads at the SNP position. This set is referred to as the “Standard Filtered” SNPs. We then used the SnpSift *filter* command with the query string “((REF=‘G’& ALT=‘A’) | (REF=‘C’ & ALT=‘T’))” to catalog the subset of the standard filtered SNPs that were either G to A, or C to T substitutions.

To detect heterozygous SNPs, we modified the above procedure for detecting homozygous SNPs. We replaced the SnpSift *filter* query string “(QUAL>=20) & (isHom(GEN[0]))& (isVariant(GEN[0]))” with “(QUAL>=20) & (isHet(GEN[0]))& (isVariant(GEN[0]))” to select the heterozygous Q20 SNPs. The low depth of reads in our genomic sequencing required a relaxed stringency for the presence of mapped reference reads at the heterozygous SNP genomic position to retain likely heterozygous positions. We accomplished this by replacing the SnpSift *filter* query string “(((DP4[0]=0) & (DP4[1]=0)) & ((DP4[2]>0)& (DP4[3]>0)))” with “((DP4[2]>0)& (DP4[3]>0))”.

### Indels detection and refinements

Indels in the variant call files were cataloged using the SnpSift *filter* command with query string “(exists INDEL)”. Homozygous and heterozygous indels with variant quality of at least 20 were retained using the *filter* command with query strings “(QUAL>=20) & isHom (GEN[0])) & (isVariant(GEN[0]))” and “ (QUAL>=20) & (isHet(GEN[0])) & (isVariant(GEN[0]))”,respectively. To eliminate homozygous and heterozygous indels from positions with strand-biased alignment we required variant calls on both strands using the *filter* query strings “(((DP4[0]=0)& (DP4[1]=0)) & ((DP4[2]>0)& (DP4[3]>0)))” and “((DP4[2]>0)& (DP4[3]>0))”, respectively.

### Detection of probable error-prone genomic positions

Following the set of variant filters described above, the standard filtered SNPs detected in the preliminary set of 570 sequenced individuals were concatenated into a single list. These alleles were sorted, by chromosome and genomic position. The number of appearances of each genomic position among the called SNPs was tallied. To the extent that EMS mutagenesis is stochastic and even in its ability to affect mutation, the same SNPs should rarely occur in more than one individual in an organism with a sufficiently large genome size. Thus, positions with a tally of two or more SNP call instances were recorded as likely error-prone positions. We refer to this set of proable error-producing positions in the sorghum reference genome (version 2.1) as the Error-Prone SNP Positions. This list of identified positions was then used to subtract the error-prone portion of the genome from the standard filtered SNP calls for each of the final 586 sequenced individuals, and included the preliminary 570 sequenced individuals. The remaining, unrepeated SNPs were unique to each line. The subtracted repeat SNPs and the unrepeated SNPs we refer to as the Replicate and Non-Replicate SNPs, respectively. We applied the same technique to the indel variant calls using the indel event starting nucleotide position of each indel as the sort item. Indels starting at positions present in more than one line were removed from all indel lists. Lists of these positions and the counts of observed SNPs in the 586 lines are provided as supplementary materials.

### Classification of SNPs and indels effect on gene function

We used the SnpEff program (version 3.1) for predicting and classifying the effects of the detected SNPs and indels on sorghum gene function. To use SnpEff, we had to create a custom SnpEff database and entry for the sorghum genome reference assembly (version 2.1). First we obtained the publicly available sorghum genome annotation and proteome files, and renamed them as “genes.gff” and “protein.fa”, respectively. These two files were installed into the subfolder for sorghum we created, in the pre-existing “data” subfolder in the SnpEff installation. We added a FASTA-formatted file containing the sorghum genome reference sequences into the pre-existing “genome” subfolder in the above “data” folder. We then edited the “snpEff.config” file to add the following entry for sorghum: “Sbicolor_v2.1_255.genome : Sorghum_bicolor”. Finally, we used the SnpEff *build* command with parameters “-gff3 -v” to construct a custom SnpEff database for sorghum. Predicted effects of the detected SNPs and indels were computed by the SnpEff *eff* command with parameter option “-c”. The output file was then postprocessed using a custom program, to retain the subset of variants predicted to have either medium or high impact on an encoded protein sequence. Finally the genome-wide distribution of indel lengths was estimated using the vcftools program (version 0.1.14) with parameter option “--hist-indel-len” (Danecek et al., 2011).

### Gene function annotation

We used the predicted protein sequences of *Sorghum bicolor* BTx623 (version 2.1), maize B73 (annotation version 5b.60) and *Arabidopsis thaliana* (TAIR release version 10) from Phytozome (version 9.1) to annotate the SNP-encoding genes. We created protein BLAST search databases for the maize and Arabidopsis sequences using the *makeblastdb* command in the BLAST suite of programs (Altschul, Gish, Miller, Myers, & Lipman, 1990; Camacho et al., 2009; version 2.2.30+), with the non-default parameters: “-input_type fasta –dbtype prot”. We aligned each sorghum protein sequence to the maize and Arabidopsis protein sequences using the *blastp* (protein BLAST) command with non-default parameters: “-evalue 1E-05 - num_threads 16 -max_target_seqs 5- outfmt 6 -seg yes”. The gene function description files for sorghum (version 2.1), maize B73 (version 5b.60) and Arabidopsis (TAIR version 10) were obtained from Phytozome and annotations were appended as additional columns to the variant call files for the affected sorghum locus along with the gene identifiers and annotations for the best maize hit and best Arabidopsis BLAST hit. The top hit for maize and Arabidopsis were appended only when BLAST e-values were less than a threshold value of E-30.

### Highly decoupled variant detection and annotation pipeline

To expedite the processing and analysis of the large-scale NGS reads data, we implemented a highly decoupled pipeline for the variant detection, filtering and annotation. The variant detection methods described above were specified as a pipelined datapath consisting of seven distinct pipeline stages, with a separate computer script representing each stage. For each sequenced sample, a custom automated script emitted separate scripts for all seven stages, along with a resource dependency graph specifying when the computer script for a particular pipeline stage should be executed on the high-performance computing system. We used the qsub job submission program with the directive “-W depend=afterok” to specify the resource dependencies among the computer scripts denoting the various pipeline stages. Using this approach greatly improved the processing and analysis throughput by leveraging the unused resource bandwidth of the computing cluster system (See Supplementary Data 21).

### Search for sequence motifs in flanking sequences of SNPs

We converted the VCF-formatted variant call files containing the non-replicate and likely EMS-induced homozygous SNPs, into the BED format using the following *awk* script command: *awk ‘{print $1“\t”$2-11“\t”$2+10}’*. This command replaces the genomic position (in the VCF format specification) with the respective start and end positions of a 21 bp window that is supported by the BED format specification. The set of flanking sequence context of all the SNP positions was saved as both FASTA and TAB delimited formats using the BEDTools *getfasta* program with parameters “-bed -fo” and “-bed -tab” respectively. Using the combination of the *srand48()* and *lrand48()* system functions in the C programming language, a subset of 10,000 sequences were randomly selected sequences with uniform distribution from the FASTA-formatted file containing the 21-nt long sequences (SNP and each flank). Using the sorghum reference genome sequences (version 2.1), we generated a third-order background Markov model using the *fasta-get-markov* utility program in the MEME ungapped motif finding software suite (Bailey et al., 2009; version 4.9.1) with non-default parameter: “-m 3”. To detect an enrichment of ungapped motifs in the sequence context of the 10,000 selected SNPs, we used the parallel version of the *meme* program for ungapped motif discovery with parameters “-dna -mod zoops -revcomp -bfile BKGRND -oc directory -maxsize 300000 -p 16 -nmotifs 50 -minw 5 -evt 0.01”. Using these parameters we limit the search results to the top 50 ungapped motifs with minimum length of 5, an e-value threshold of 0.01, and the profiling evaluated both the original sequences and their reverse complements.

### Analysis of the DNA sequence context of the entire set of probable EMS-induced mutations

The FASTA-formatted file containing the extracted DNA sequence contexts of all the likely EMS-induced SNPs was partitioned into two files, based on whether the SNP reference base was a G or C nucleotide. For each of the 20 positions surrounding the SNP, we tabulated the number of occurrences for all the four DNA nucleotide types in each sequence. The tabulation was performed separately for each of the above two files. Next we repeated the process using the same sample sizes but with two files containing 21nt sequence contexts of randomly selected G and C reference base positions in the sorghum genome. For each DNA nucleotide type, and at each of the 20 positions, we computed the percentage changes (deviations) in the number of occurrences from the random dataset to the EMS-induced SNP dataset. We analyzed the tabulations of the pairs of datasets for the G and C reference bases cases, separately.

### Detection of paralogs of the sequence contexts of error-prone SNP positions

Due to the sorghum genome complexity, we determined that around 50 flanking bases is sufficient to distinguish the sequence context of a specified genomic position from another. We hence extracted the 51nt sequence context for the detected error-prone SNP positions. These were aligned to the sorghum reference genome sequences in a BLAST-indexed database using the *blastn* program in the BLAST software suite (version 2.2.30+) with parameters options: “-evalue 1.0E-10 -num_threads 16 -outfmt 0 -ungapped -dust no -word_size 11”. The BLAST e-value threshold was selected based on initial preliminary runs in order to exclude a high percentage of spurious matches in the alignment results. The BLAST unngapped alignment output results were initially parsed to retain only the subset of alignment results, which consists of global alignments. The stringency of the filtering was subsequently increased to only retain global alignments containing a single substitution, and coinciding with the 26^th^ position (error-prone SNP position). The alignments and the single nucleotide substitution base were then catalogued. For each of the error-prone SNP positions, we compared the DNA base in the single nucleotide substitution in the BLAST alignments with the false positive SNPs calls at the error-prone reference position, and kept track of matching and mismatching bases. We repeated this procedure for two other datasets, with each consisting separately of 58,000 randomly selected genomic positions, and likely EMS-induced homozygous SNPs. We tabulated and compared the BLAST results for the three datasets.

### Annotation of likely adverse effect of missense mutations using SIFT

We followed the recommended steps for the setup and application of the SIFT program for predicting the effect of missense mutations on protein function (Kumar, Henikoff, & Ng, 2009; version 5.2.2). We downloaded the UniRef90 protein database (Consortium, 2014; Suzek, Huang, McGarvey, Mazumder, & Wu, 2007; release 2015_03) from The European Bioinformatics Institute website (http://www.ebi.ac.uk/uniprot; retrieved in March, 2015). The FASTA-formatted sequences were then formatted into a protein BLAST database using the *formatdb* command in the BLAST package with parameters: “-i uniref90.fasta -p T –n uniref90”. The SnpEff annotated VCF-formatted files were parsed to extract the subset of likely EMS-induced which were predicted to cause missense mutation. The output file consisted the amino acid changes, their positions in the protein sequences and the genes that encode them. Using this table, we generated an input file for each sorghum gene transcript with at least one missense EMS-induced SNP. The file for each gene transcript contained all the amino acid changes detected for that gene in all 586 sequenced individuals. The amino acid sequence of each affected gene transcript was downloaded from Phytozome and stored in separate FASTA-formatted files. The individual sorghum protein sequences, UniRef90 BLAST database and amino acid changes file for each sorghum gene transcripts were used as input to the *SIFT_for_submitting_fasta_seq.csh* C-shell utility script that was provided in the SIFT program to predict the effect of missense amino acid changes on protein function. For SIFT prediction, we specified a median conservation value of 2.75 and a SIFT score threshold of 0.05. The *seqs_chosen_via_median_info.csh* script in the SIFT package was modified to increase the CPU time limit from 30 to 60 minutes (“limit cputime 60m”) and the *psiblast* alignment program was executed in the multithreaded mode (“-num_threads 16”) instead of the default single threaded CPU mode. This specified the parallel processing for each missense SNP-encoding gene transcript and significantly reduced analysis time. A mutation was predicted to be deleterious if the computed SIFT score is lower than the threshold. We parsed the output results of the SIFT program to retain amino acid changes with predicted “DELETERIOUS” and “TOLERATED“ designations. These designations were appended to the corresponding amino change columns in the EMS-induced SNPs annotation file.

## RESULTS

### Generation of a *Sorghum bicolor* BTx623 EMS population

A population of BTx623 sorghum was treated with 45 mM EMS to induce mutations. Taking advantage of the primarily selfing habit of sorghum, plants were open pollinated and ~12,000 lines were collected. Over the following two summers, M_1_ seed from 7,304 of these were planted and one individual from each of 586 families that were advanced to the M_3_ generation was used to isolate DNA for sequencing. Each of these selected plants was also self-fertilized by controlled pollination and M_4_ seeds reserved for stock center deposit, phenotypic assessment, and future research.

### NGS sequencing and reads mapping

A total of 29,972,216,082 Illumina next-generation sequencing paired-end reads were generated from 586 EMS-mutagenized sorghum BTx623 individuals (Table 1). Of these reads, 29,651,940,529 (99%) aligned to the sorghum BTx623 reference genome (Paterson et al., 2009). From the alignment, the estimated median paired-end insert size was 527 base pairs. The average genome coverage of the alignments was 7X. These data were used to identify SNPs likely to be the result of the EMS treatment. This sequencing depth is lower than the 20X coverage depth often described as minimum for accurate SNP calling (Flibotte et al., 2010; James et al., 2013; Liu, McCormack, & Sheen, 2012). We sought to control for false positive SNP calls by considering the data from across the experiment as an estimate of error, and processing the data as follows.

**Table 1.** Summary sequencing and alignment statistics for the 586 EMS-mutagenized sorghum BTx623 individuals

|  | **Total** | **Mean** |
| --- | --- | --- |
| **Reads** | 29,972,216,082 | 51,147,126 |
| **Mapped Reads** | 29,651,940,529 | 50,600,581 |
| **Properly Paired Reads** | 29,288,740,978 | 49,980,787 |
| **Median Insert Size** | - | 526 |
| **Genome Coverage** | - | 7 |

### Standard SNPs detection and filtering

To detect the high quality EMS-induced SNPs from the nearly 30 billion sequencing reads, we implemented a highly decoupled SNPs prediction and annotation pipeline (See Methods). Initially, we detected 31,178,985 putative SNPs in the 586 EMS-mutagenized individuals using the SAMtools package (H. Li et al., 2009). The default criteria for SNP calling resulted in an unreasonably high number of positives: a polymorphism for every ~14kb of genome with 53,206 SNP sites per individual (Table 2). The removal of SNPs of low quality, repetitive origin or low reads coverage reduced this number by only ~10% to 28,472,909 “Repeats-Filtered” SNPs (Table 2). These M_3_ self-pollinated plants should predominantly include homozygous, induced mutations. To retrieve the homozygous subset of SNPs, we filtered the SNPs to retain only those positions predicted to be homozygous and having a SNP quality score of at least 20, which we refer to as the homozygous “Q20” SNPs. This filter was much more effective and only 24% (6,906,590) of the SNPs were retained, with an average number of 11,786 homozygous Q20 SNPs per individual (Table 2). Previous studies show strand-biased SNPs tend to be a source of false positive SNP detections (DePristo et al., 2011). We removed SNPs derived from reads mapping to only one strand of the genome, leaving a total of 5,399,493 high-quality SNPs, and an average of 9,214 per individual (Table 2; Supplementary Data 1). We designate this set as our “Standard Filtered” homozygous SNPs. We applied the same filtering steps to the heterozygous SNP calls. This retained 7,444,547 standard filtered heterozygous SNPs, and an average of 12,704 per individual (Supplementary Table 1; Supplementary Data 1).

**Table 2.**
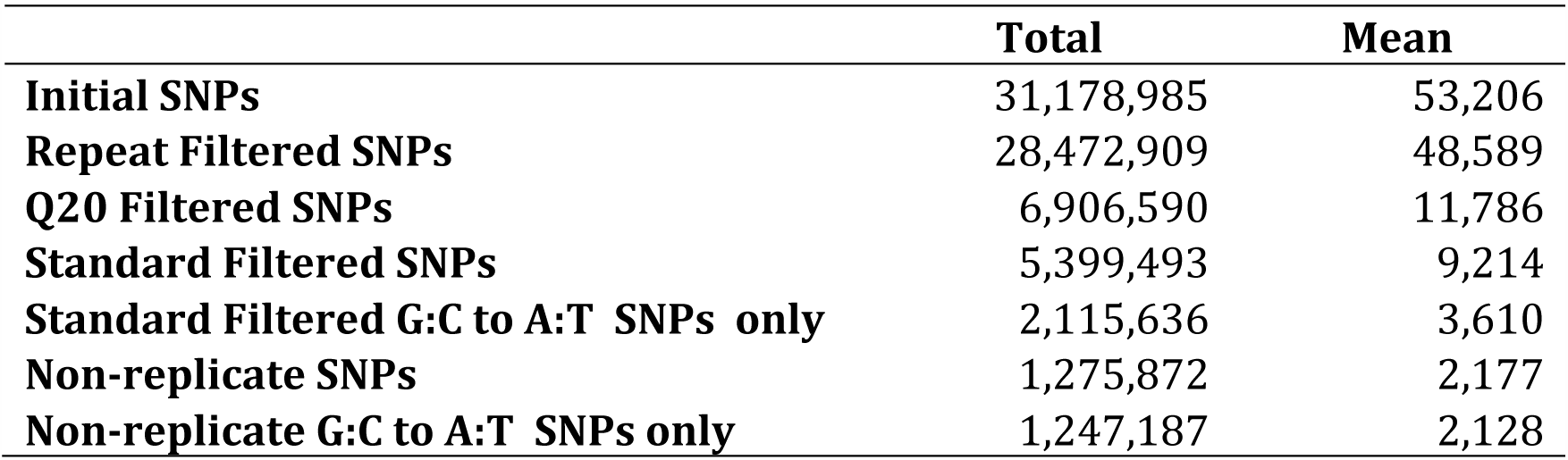
Summary statistics for the detection of homozygous single nucleotide polymorphisms in the 586 EMS-mutagenized sorghum BTx623 individuals

### Prediction of error-prone SNP positions

Because pollen and seed contamination might result in the mistaken resequencing of the same line twice, we compared the SNP calls between lines. This analysis identified 16 lines with greater than 68% overlap between the called SNPs and patterns of SNP coincidence that were consistent with pollen or seed contamination. For additional analysis, we therefore focused on the 570 lines without siblings. For these 570 lines, the SNPs detection and standard filtering retained a total of 5,244,176 standard filtered homozygous SNPs produced at 1,290,635 distinct genomic positions. Surprisingly, 4,011,413 of the 5,244,176 SNPs called were predicted repeatedly from only 57,872 distinct genomic positions. This means that roughly 4% of the distinct locations produced more than 76% of the SNP calls in the sequence analysis of these 570 individuals (Figure 1; Supplementary Table 2). For example, there were a total of 1,085 positions in the sorghum genome, where the same SNP was detected in between 451 and 500 EMS-treated sorghum individuals. This group of SNPs accounted for 515,902 redundant SNP detections (see Supplementary Table 2). We refer to these repeatedly predicted SNPs as the Replicate SNPs. These positions are not random poor base calls. For example, for nearly all (57,802 positions) of these Replicate SNPs, the predicted change was the same for all lines (Supplementary Data 2). For example, in 567 lines a C to G substitution was predicted at the genomic position 37,580,764 on chromosome 5. This pointed to some structural basis for the repeated observations. Similarly, of the 7,238,377 standard filtered heterozygous SNPs, 6,775,202 SNPs were derived from replicate SNPs at only 71,294 positions. Thus, roughly 13% of the 534,469 distinct genomic positions produced replicate heterozygous SNPs accounting for 94% of the heterozygous SNP predictions in the 570 sorghum individuals (Figure 1).

**Figure 1.**
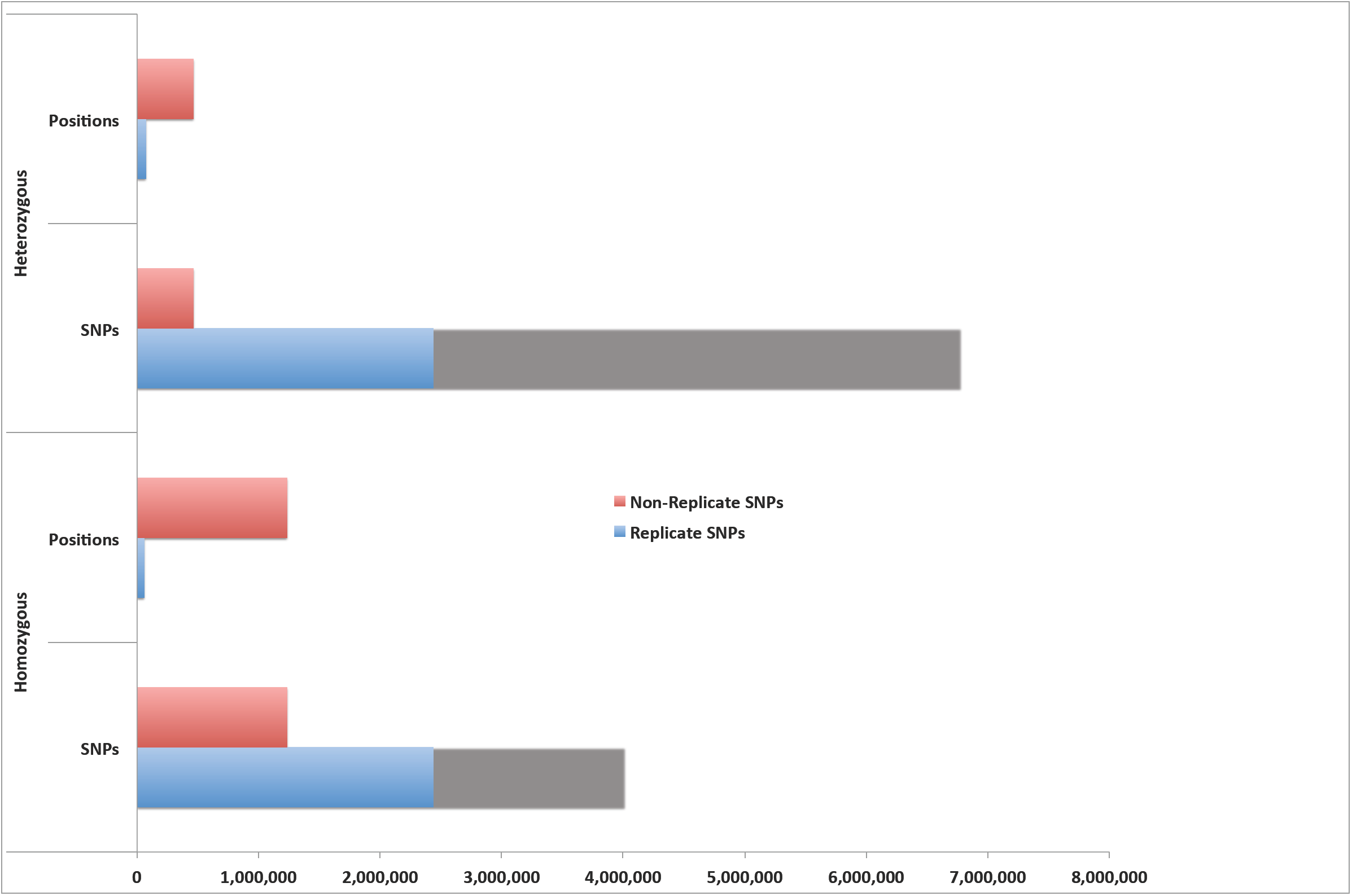
Detection of the subset of likely EMS-induced SNPs in the standard filtered SNPs. Over 76% of the standard-filtered homozygous SNPs were predicted repeatedly from only 4% of the total distinct homozygous SNP positions in 570 sorghum individuals. For the standard-filtered heterozygous SNPs 94% were predicted repeatedly from only 13% of the distinct heterozygous SNP positions.

EMS chemical mutagenesis results primarily in G:C to A:T mutations at random genomic positions (Loveless, 1958; Loveless, 1969; Sega, 1984; Shrivastav, Li, & Essigmann, 2010; Waugh et al., 2006). If the replicate SNPs are largely the result of hotspots for EMS-mutagenesis, one would expect more than 90% of SNP calls to be G:C to A:T mutations at these positions. However, only 39% (2,050,166 SNPs) of the standard filtered homozygous SNP predictions were G:C to A:T substitutions (Supplementary Figure 1; Supplementary Table 3). Similarly, only 36% (2,588,504 SNPs) of the heterozygous SNPs were G:C to A:T substitutions. With less than 40% of the retained high quality standard filtered SNPs showing signatures of EMS-derived mutagenesis, it is more likely that a significant proportion of the 5,244,176 homozygous and 7,238,377 heterozygous SNPs are false positives. Since *Sorghum bicolor* BTx623 has a sufficiently large genome size (Arumuganathan & Earle, 1991; Peterson et al., 2002; Price et al., 2005; Paterson et al., 2009) with the *in silico* genome reconstruction (version 2.1) containing nearly 727 megabases (Paterson et al., 2009), and given that all the 570 samples were treated independently with EMS, the probability of detecting an EMS-induced SNP at the same genomic position in multiple sorghum individuals is low. We propose that the predicted Replicate SNPs are most likely to be false positives and therefore these were subtracted. After subtraction of the Replicate SNPs, those positions that were unique to an individual remain, and we designate these as Non-replicate SNPs. Remarkably, consistent with the experimentally determined effect of EMS mutagenesis, 98% (1,205,140) of the retained, non-replicate, homozygous SNPs were G:C to A:T transitions (Supplementary Figure 2; Supplementary Table 3), The second and third largest substitution classes were A:T to G:C transitions and A:T to T:A substitutions, accounting for 0.7% (9,066) and 0.5% (6,194) of the SNPs, respectively. Similarly, in the case of the heterozygous standard filtered SNPs, after removal of the replicate SNPs, the remaining, non-replicate SNPs were 94% (435,414) G:C to A:T transitions (Supplementary Figure 2; Supplementary Table 3).

The detected error-prone SNP positions in the 570 unique lines were used to mask likely false positives from the standard filtered SNPs called for each of the 586 lines. The additional 16 sorghum individuals with overlapping genomic positions were excluded from the masking set, so as not to remove true EMS-induced SNPs that were due to contaminated pedigrees. Cumulatively, 1,275,872 (24%) of the 5,399,493 standard filtered homozygous SNPs were non-replicate SNPs in the 586 sorghum individuals (Table 2; Supplementary Data 3), and 98% (1,247,187) were G:C to A:T transitions (Figure 2; Supplementary Table 4). Protein-coding gene loci comprised 246,062 (19%) of the likely homozygous EMS-induced SNPs and over 99% (243,775) were G:C to A:T mutations (Figure 2). Similarly, 477,531 non-replicate heterozygous SNPs (6%) were retained in the 586 lines, after removal of replicates in the 7,444,547 heterozygous standard filtered SNPs (Figure 2; Supplementary Tables 1 and 4; Supplementary Data 4). Using the final combined 1, 753,403 non-replicate SNPs, we examined the level of SNP homozygosity in each mutagenized individual by comparing the number of *in silico* detected homozygous and heterozygous SNPs in each sample. The ratio of homozygous to heterozygous SNPs for the population was 2.7:1. Sorghum is a diploid organism so homozygous variants were present on both copies of each chromosome, while the heterozygous variants were present on one only. Hence, the observed ratio of non-replicate SNPs that were homozygous to SNPs that were heterozygous SNP count ratio is 5.4:1. A total of 515 individuals (88%) were homozygous for at least 60% of their SNPs, and only 21 samples were at most 20% homozygous (Figure 3; Supplementary Data 5). Overall, we predicted an average of 2,177 homozygous and 815 heterozygous non-replicate SNPs per individual, and estimated an EMS mutation frequency of about one SNP per 243 kilobases (with one homozygous SNP per 334 kilobases).

**Figure 2.**
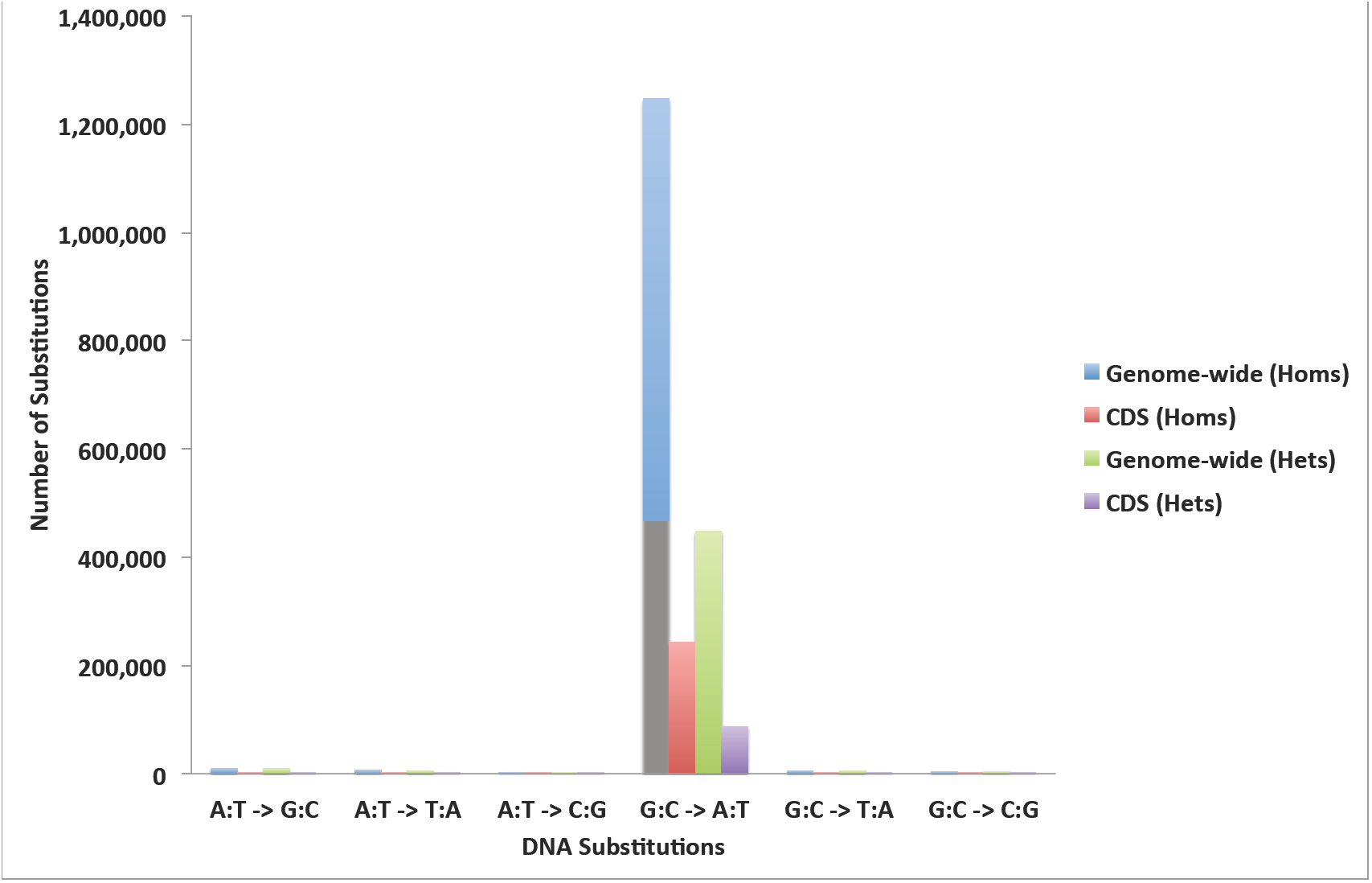
The mutation spectrum and frequency distribution of the probable EMS-induced homozygous and heterozygous mutations in the whole genomes and protein-coding regions of the 586 resequenced sorghum BTx623 individuals.

**Figure 3.**
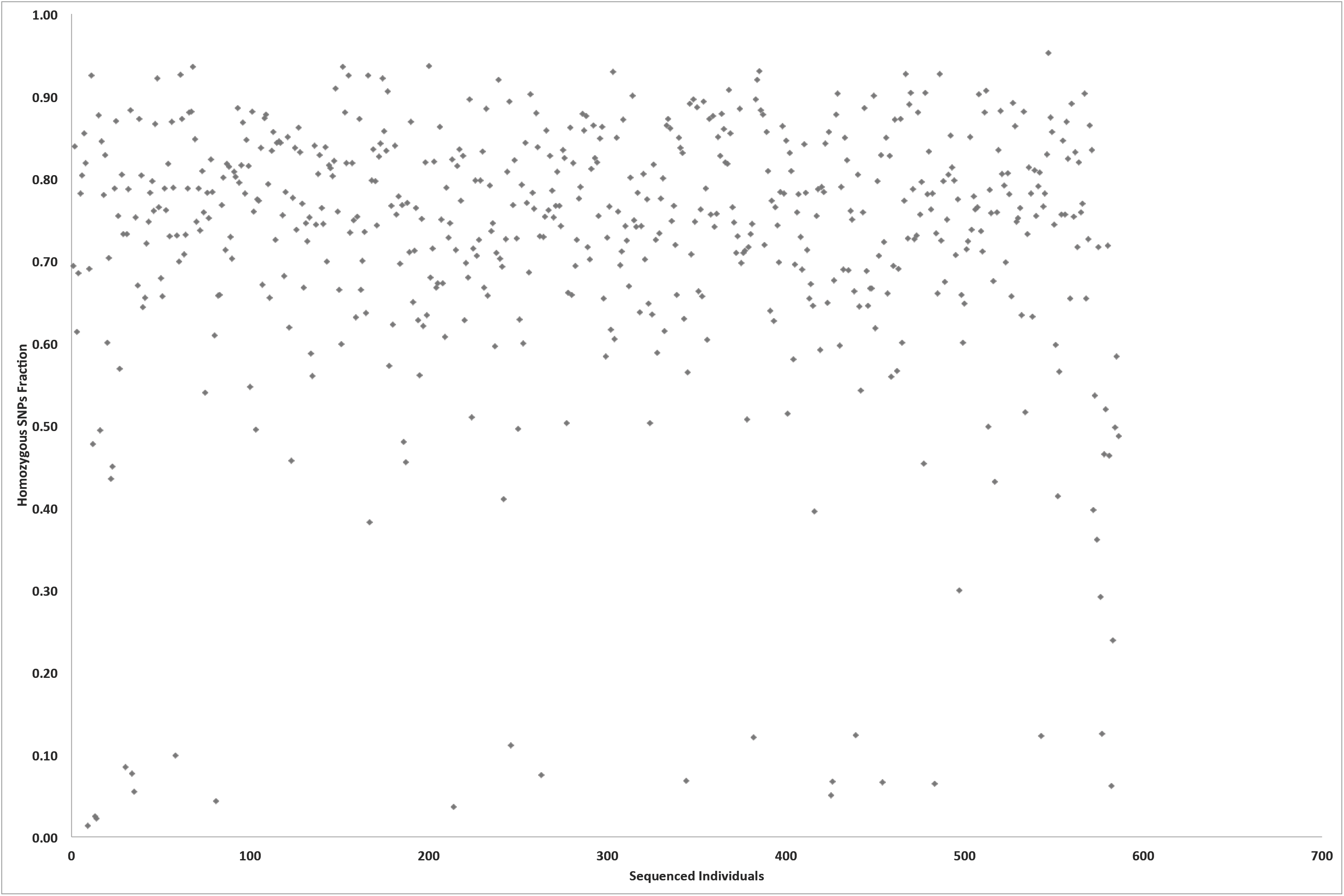
The level of SNPs homozygosity in the genomes of the 586 EMS-treated sorghum BTx623 individuals.

### Functional classification of SNPs and their impacts

The majority of the SNP calls were outside predicted protein-coding regions. Intergenic regions and introns contained 1,032,427 (81%) and 109,144 SNPs (9%) of the non-replicate homozygous SNPs, respectively (Table 3; Supplementary Table 5; Supplementary Data 6 and 7). The remaining 134,301 SNP calls were in protein-coding sequences, with 60,442 effecting changes in protein-coding capacity in 23,866 distinct sorghum genes. Homozygous mutations of high impact (loss of reading frame) were predicted for 4,035 SNPs disrupting 3,637 distinct genes. These mutations included 2,880 nonsense alleles and 1,043 mutations removing splice sites. A total of 56,407 non-replicate, homozygous SNPs were predicted to cause nonsynonymous amino acid substitutions in 23,198 distinct sorghum genes. In addition, we detected five lost stop codons associated with non-replicate SNP changes. These five mutations were not G:C to A:T mutations, as no such mutations can induce a stop codon loss, which may explain the relatively small number relative to the 145 start codon losses. The stop codon losses included a single homozygous and four heterozygous mutations. Synonymous changes from one stop codon to another were associated with G:C to A:T mutations and we detected 148 synonymous (silent) stop codon substitutions consisting of 114 homozygous and 34 heterozygous mutations. The silent stop lost mutations include 92 TGA and 56 TGA codon changes to TAA codons (Supplementary Table 6; Supplementary Data 8). Mutations causing loss of start codons included 111 homozygous and 34 heterozygous changes (Table 3; Supplementary Table 5). Intergenic regions and introns contained 387,570 (81%) and 42,357 (9%) of the 477,531 non-replicate heterozygous SNPs, respectively. The heterozygous SNPs also included 1,813 high impact and 19,800 non-synonymous mutations (Table 3; Supplementary Data 9 and 10). Taken together, counting mutations in exons, splice sites and untranslated regions we detected 188,371 alleles in 31,163 (94%) sorghum genes, including 139,600 homozygous alleles in 30,167 genes (Figure 4). Within protein-coding sequences we also predicted 79,765 nonsynonymous and 46,234 synonymous mutations, as expected based on codon degeneracy. On average, we predicted seven high impact (e.g. nonsense mutation or loss of splice site) and 96 amino acid substitution alleles per individual. The average individual also encoded three high impact and 34 moderate impact non-replicate heterozygous SNPs per individual. We estimated an average of 5.7 alleles per sorghum gene.

**Table 3.**
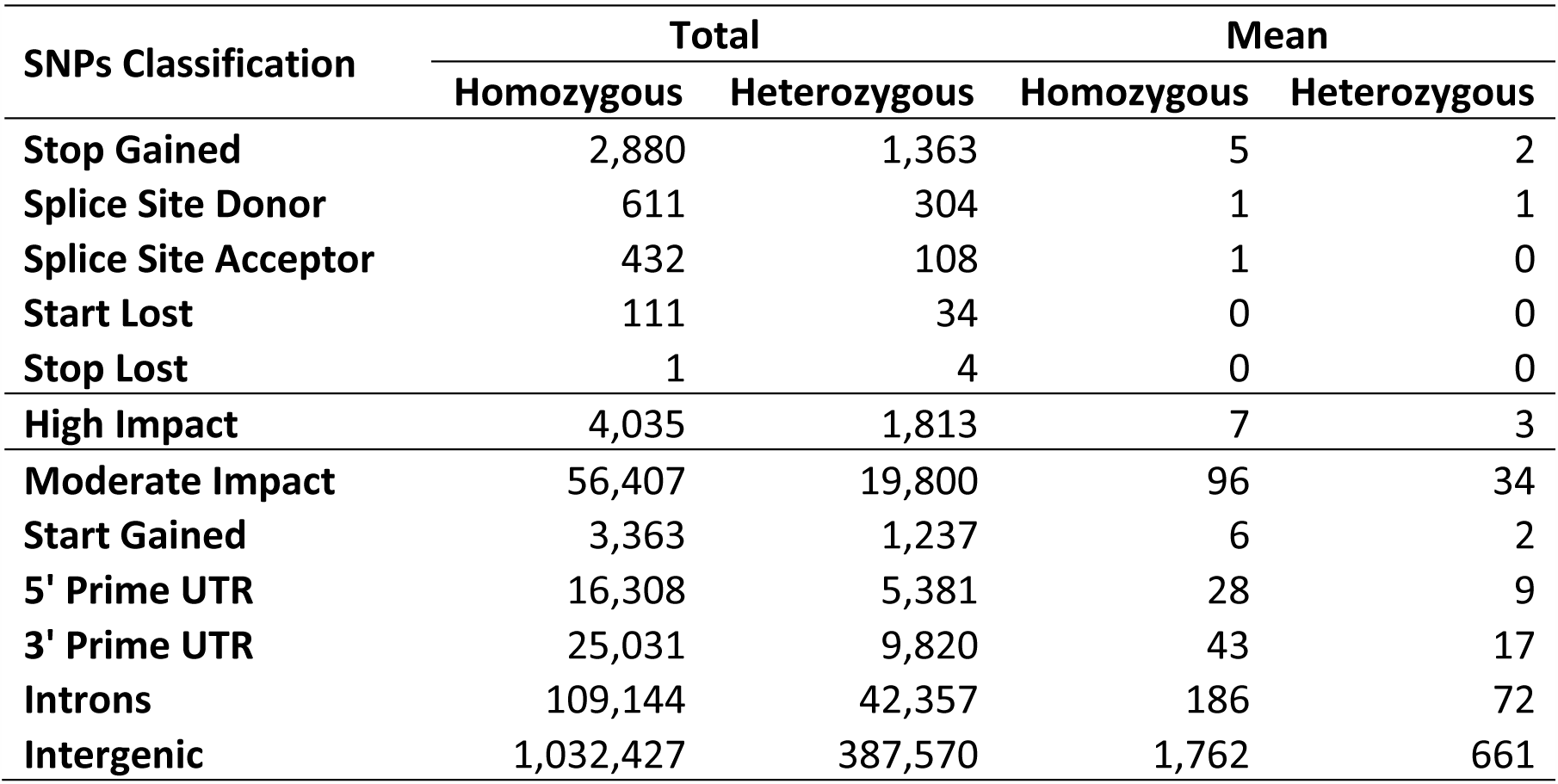
Classification of the predicted EMS-induced SNPs and their effects in the 586 EMS-mutagenized sorghum BTx623 individuals.

| **SNPs Classification** | **Total** |  | **Mean** |  |
| --- | --- | --- | --- | --- |
|  | **Homozygous** | **Heterozygous** | **Homozygous** | **Heterozygous** |
| **Stop Gained** | 2,880 | 1,363 | 5 | 2 |
| **Splice Site Donor** | 611 | 304 | 1 | 1 |
| **Splice Site Acceptor** | 432 | 108 | 1 | 0 |
| **Start Lost** | 111 | 34 | 0 | 0 |
| **Stop Lost** | 1 | 4 | 0 | 0 |
| **High Impact** | 4,035 | 1,813 | 7 | 3 |
| **Moderate Impact** | 56,407 | 19,800 | 96 | 34 |
| **Start Gained** | 3,363 | 1,237 | 6 | 2 |
| **5' Prime UTR** | 16,308 | 5,381 | 28 | 9 |
| **3' Prime UTR** | 25,031 | 9,820 | 43 | 17 |
| **Introns** | 109,144 | 42,357 | 186 | 72 |
| **Intergenic** | 1,032,427 | 387,570 | 1,762 | 661 |

**Figure 4.**
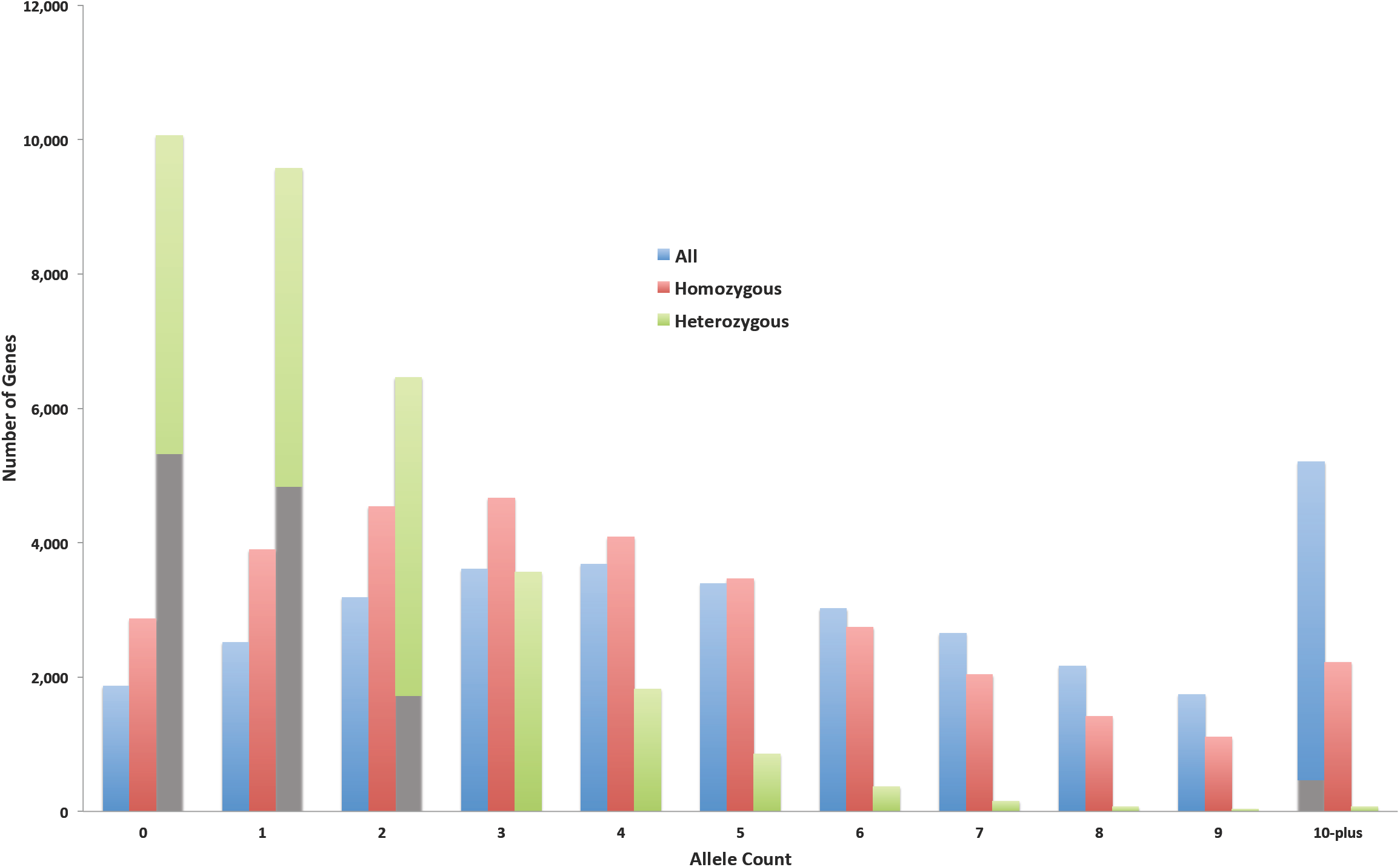
Frequency distribution of allele counts for all genes impacted by likely EMS-induced mutations in protein-coding exons, splice sites and untranslated regions of the sequenced 586 sorghum BTx623 individuals.

### Prediction of insertion and deletion polymorphisms

We initially detected 3,312,791 insertion/deletion (indel) polymorphisms in the 586 sorghum lines and the standard filtering retained 1,989,078 indels (Supplementary Tables 7 and 8; Supplementary Data 11 and 12). Replicate Indels were detected and filtered out by identifying all indels that start at the same genomic position in multiple individuals. Less than 1% (17,760) of the remaining indels were retained after this process, indicating an unacceptably high rate of false positives. The average length of the retained indels was 2.6 bp with a remarkably skewed distribution (Supplementary Figure 3). Among the non-replicate indels,77% were up to 2 bp in length and 97% of the indels were 10 bases or shorter. A total of 277 and 180 indels were predicted to have high and moderate impact on gene function, respectively. One deletion, in line SbEMS 0456-2, disrupts the sorghum ortholog, Sobic.003G386900, of the *Monoculm1* gene from rice, LOC_Os05g42130. The phenotype of this line is reduced height and a complete absence of tillers. Twenty-two indel variations affected splice sites and 1,094 occurred in untranslated regions of sorghum protein-coding genes. Another 14,341 and 1,947 indels occurred in intergenic and intronic regions, respectively.

### Predicting the impact of missense SNPs on protein function

Over 93% (68,379) of the 73,346 homozygous non-synonymous EMS-induced SNP were missense mutations. These affected the coding sequences of 27,943 transcripts encoded by 23,227 distinct sorghum genes. To provide additional guidance for later data re-use, we used the SIFT program in conjunction with the 32,168,296 sequences in the high quality UniRef90 protein database to predict the likely impact of these SNPs on their encoded proteins. We were able to predict the impact of 56,289 missense SNPs. Of these, 25,013 (44%), affecting 12,840 genes, returned a SIFT score <0.05 and likely encode changes deleterious to protein function. The remaining 31,276 SNPs were more likely to be tolerated mutations in 14,559 genes (Supplementary Data 13). Similarly, 24,191 (92%) heterozygous missense mutations were found in 16,118 transcripts encoded by 13,340 (40%) distinct sorghum genes. SIFT analysis predicted the impact of 21,183 of these heterozygous mutations and 9,659 (46%) of them were predicted to be deleterious changes. The remaining 11,524 heterozygous SNPs affecting 7,417 genes were classified as likely tolerated mutations (Supplementary Data 14). The detailed summary statistics on the sequencing, mapping, SNPs prediction, filtering, annotation and classification for the homozygous and heterozygous SNPs, as well as the indel polymorphisms are included as supplemental information (see Supplementary Data 15). The detailed classification of the medium or high impact SNPs and indels and annotations for the affected genes in all 586 sequenced lines are included as supplementary information (See Supplementary Data 16, 17, and 18).

### Classification of genes with no detected EMS-induced mutations

The sorghum genome annotation includes 39,441 transcripts derived from 33,032 distinct protein-coding genes. We detected SNPs in 89% of these genes (35,221 transcripts and 29,296 genes). Using the agriGO gene ontology web server (Du, Zhou, Ling, Zhang, & Su, 2010), we analyzed the remaining 3,736 genes (4,220 transcripts) lacking SNPs and found an overrepresentation of genes with biological functions in photosynthesis (4.98e-08), RNA translation (1.15e-12), generation of precursor metabolites and energy (5.88e-10), and response to wounding (2.99e-06) (Supplementary Figure 4). We next examined what attributes of a gene predict the number of mutations observed in our EMS mutagenesis. As EMS mutagenesis results predominantly in G:C to A:T substitution, we tabulated the GC-content and number of detected SNPs for all the transcripts. The GC-content of sorghum transcripts with and without SNPs was significantly different (p-value of 3.17e-32); transcripts with no SNPs had a mean GC-content of 54% whereas transcripts containing SNPs (mean = 4.6) had a mean GC-content of 56%. We found no strong transcriptome-wide linear correlation between GC-content and the number of SNPs (r^2^=0.00279; Figure 5A). Unsurprisingly, substituting GC-content (di-nucleotide percentage calculation) with G+C-count, we found the number of SNPs was linear correlated with G+C count (Figure 5B; r^2^=0.61069; Supplementary Figures 5 and 6). Transcripts containing SNPs had mean G- and C- counts of 361.1 and 340.4 respectively, while the mean count of Gs and Cs for transcripts with zero SNPs were 133 and 128.2 respectively indicating that genes without SNPs were enriched for genes that represented a smaller mutational target (Supplementary Data 19). Consistent with this, we found a transcriptome-wide linear correlation between the number of SNPs and the length of a transcript(r^2^=0.57184; Figure 5C). While the mean length of SNP-containing transcripts was 1,295, the mean length of SNP-lacking transcripts was only 480. Hence in general, transcript length and the number of G (or C) nucleotides in a transcript impact the likelihood of an EMS-induced mutation. Further mutagenesis to saturate the sorghum genome may benefit from both a greater population size and a diversity of mutagens.

**Figure 5.**
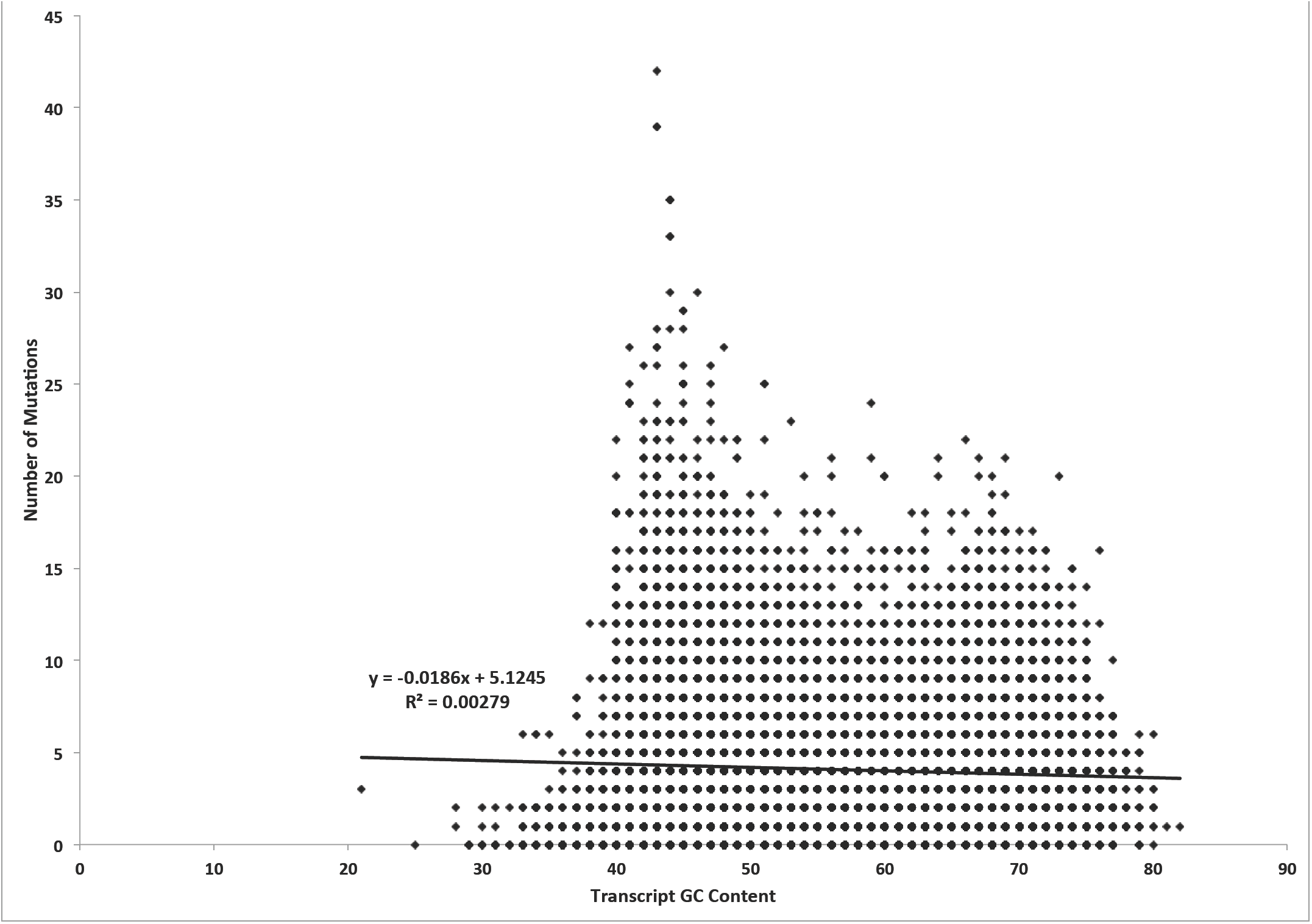
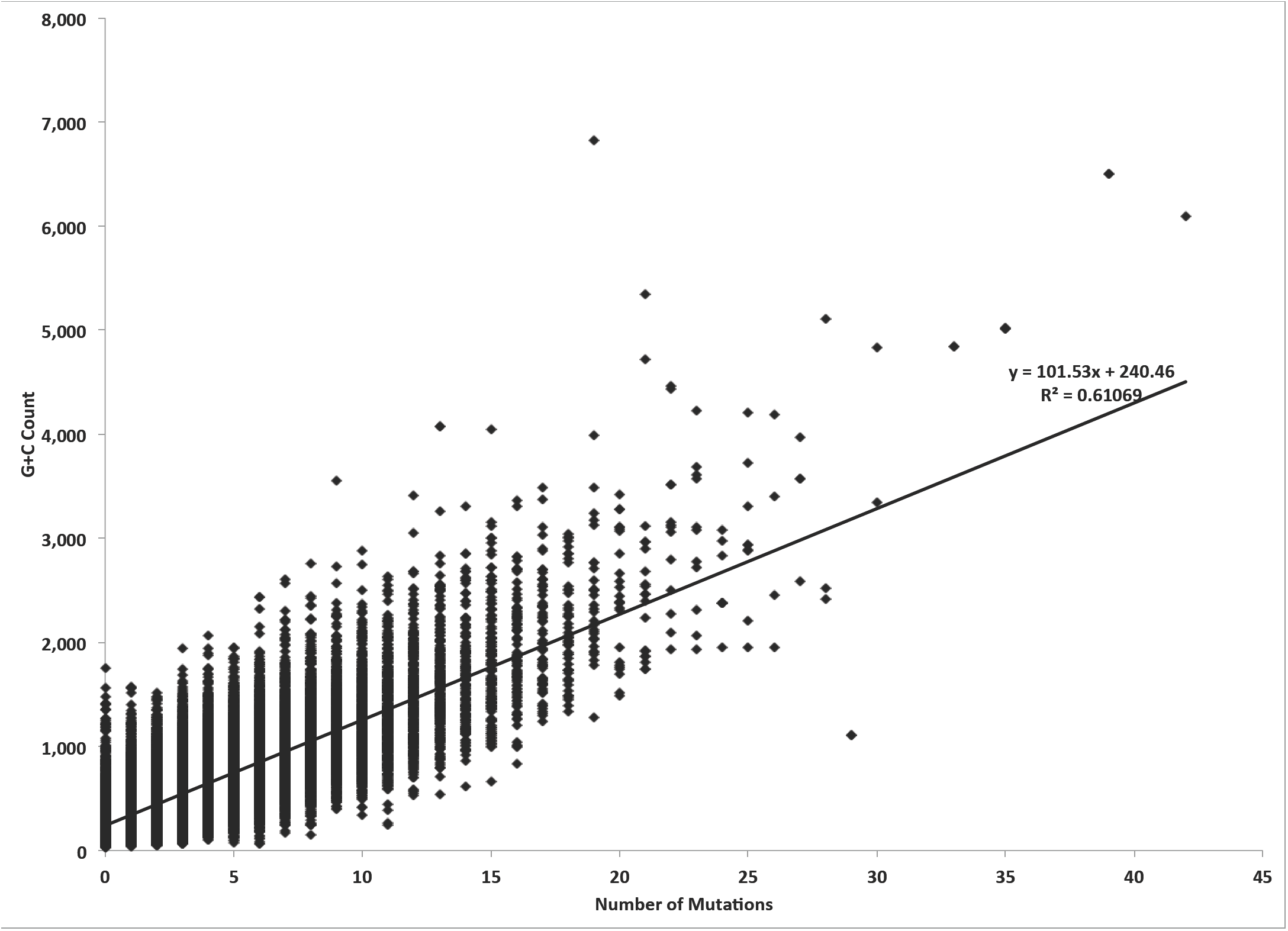
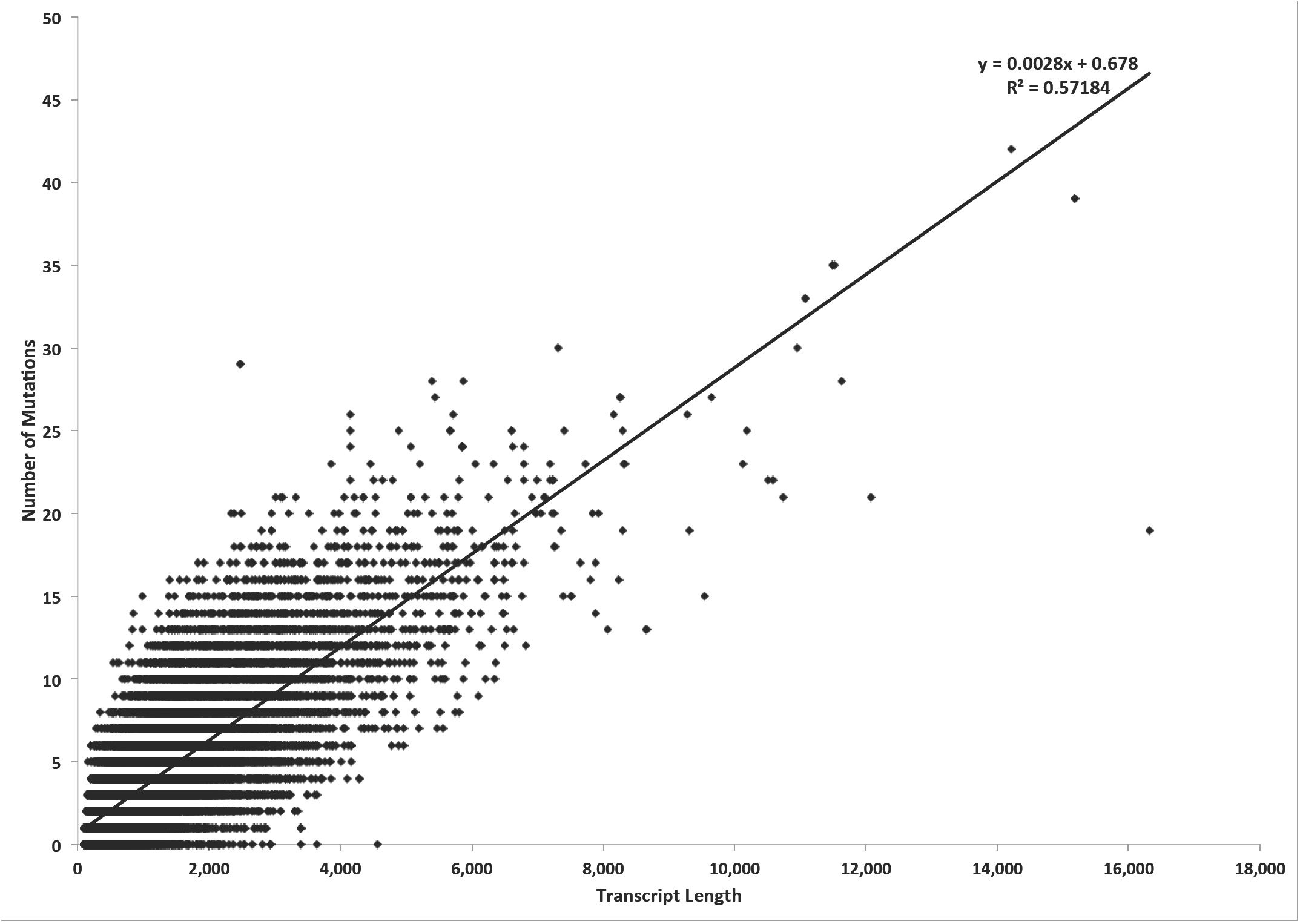
The examination of attributes that impact the number of predicted EMS-induced mutations in a gene transcript. (a) Scatterplot shows no direct linear correlation between mutation count and GC-content.(b)Scatterplot depicting a linear correlation between mutation count and G+C-count.(c) Linear correlation between mutation count and transcript sequence length.

### Sequence context of EMS-induced SNPs

We examined the sequence context of the EMS-induced SNPs for conserved flanking sequence motifs. These may enhance or promote mutagenesis and further bias the induction of mutations in the genome. Using the MEME ungapped motif search program, we found no single fixed length uniform motif associated with the 10,000 randomly selected SNPs. However, we observed an enrichment for motifs that were present in clusters of SNPs. The most significant motif (p-value of 5.7e-304) was a 16nt long GC-rich sequence, which is present in a cluster of only 834 SNPs out of the 10,000 homozygous SNPs (Supplementary Figure 7). In the absence of uniform flanking motifs, we compared the available 21nt sequence context of the 1,247,187 homozygous EMS-induced G:C to A:T SNPs, with an equal number of randomly selected G and C genomic sites from the sorghum genome. We performed separate analyses for the 608,485 and 606,560 mutated reference G and C bases respectively. Compared to the flanking sequences of the randomly selected G reference bases, we observed significant sequence changes in the single base immediately upstream (position −1), and two bases immediately downstream of the mutated genomic position (Figure 6). In the first base position immediately upstream of the mutated central G position, we observed a 28% increase (99,332 versus 127,287) in the count of Cs, and a 17% decrease (177,307 versus 146,321) in the count of A bases. Intriguingly the upsurge in the C count occurred at the position immediately preceding the mutated G, implying a greater proportion of CG dinucleotides among the mutated bases. These dinucleotides are underrepresented (3.6%) in the sorghum genome (version 2.1), and are the sites of DNA methylation. Methylated cytosines are known to spontaneously deaminate to thymidine. For this reason the G to A mutational impact of EMS, which would pair perfectly with thymidine, may be more readily affected at these locations and go undetected by repair processes. At the first downstream position (+1) we observe a 32% increase (141,485 versus 186,396) in the count of Cs and a 26% decrease (144,587 versus 106,292) in the count of Gs. At the second downstream position (+2) we observe a 38% increase (139,477 versus 192,222) in the G count and an 18% decrease (133,840 versus 109,737) in the count of Cs. The reverse complement of the results of the analysis for the mutated C reference bases yields the same pattern (Supplementary Figure 8). Based on the nucleotide counts and the percentage changes at each flanking position we observe an overrepresentation of a 5‘-C**G**CG-3’ motif (with the mutagenized G position showed in bold) and an underrepresentation of 5’-AHGM-3’ (where the ambiguous DNA codes H={A,C,T} and M={A,C} respectively). In comparison to the genome the CG dinucleotide content was 57% higher in the 21nt sequence context of the mutated positions (3.4% versus 5.3%) and the 5’-C**G**CG-3’ motif is also clearly delineated in the positions 6 to 9 in the MEME motif results.

**Figure 6.**
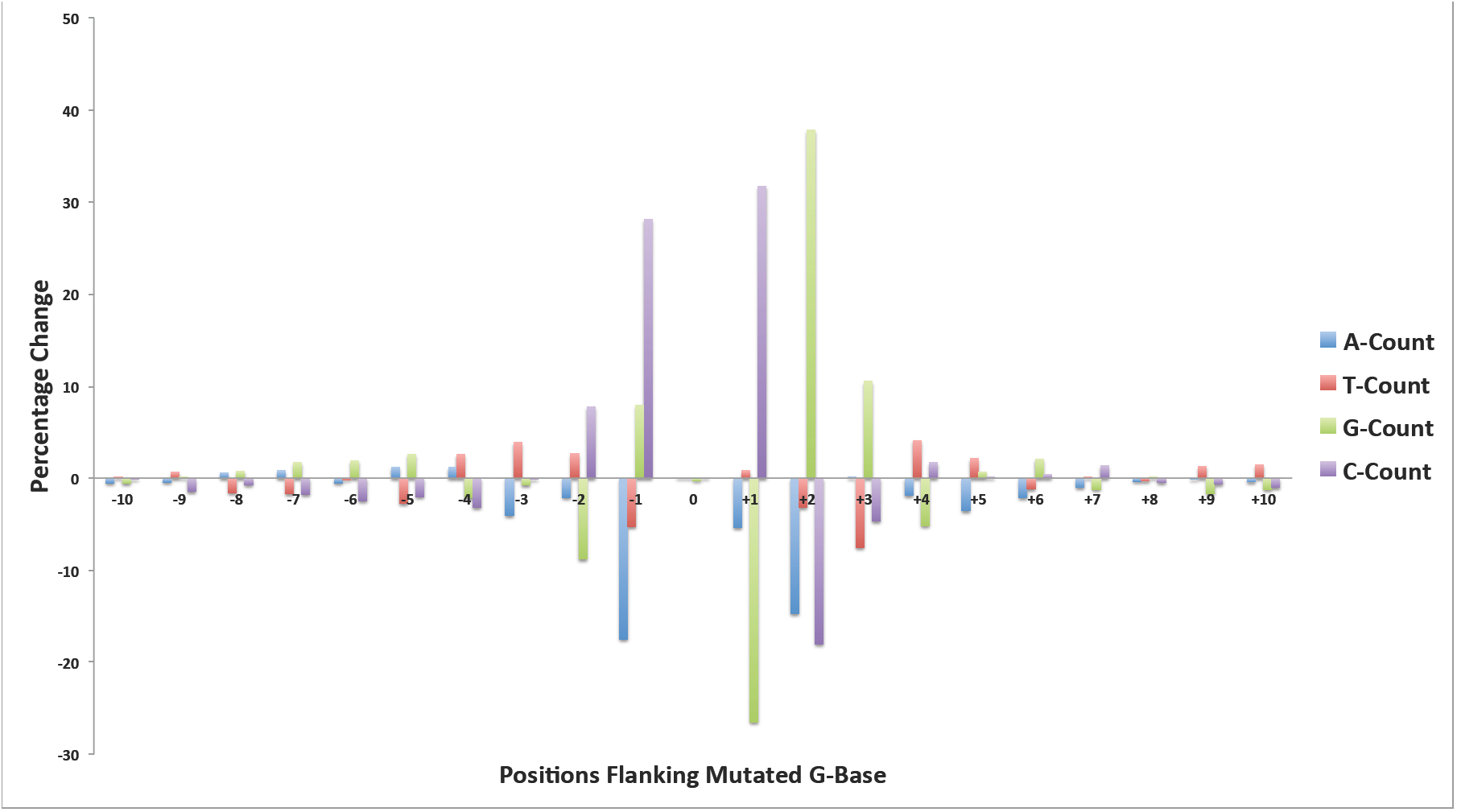
The 21nt sequence contexts comparison for the mutated G-residue DNA sites in the EMS-induced SNPs and randomly selected positions in the sorghum reference genome. The percentage change in the nucleotide frequency for all DNA base types in all DNA positions shows an overrepresentation of a 5’-C**G**CG-3’ motif. The mutated G-residue is shown in bold.

### Sequence context of the error-prone SNP Positions

We examined the contribution of paralogy to the landscape of error-prone SNP positions. We selected similar numbers of error-prone SNPs, randomly selected genomic positions, and randomly selected homozygous non-replicate EMS-induced SNPs. We then obtained the 51nt genomic sequence contexts of these positions, removing from the analyzed sets any positions that did not have unambiguous and complete sequence data for the entire 51nt. We performed ungapped global alignment using BLAST. We obtained BLAST global alignment results for 57,404, 55,085 and 57,994 of the error-prone SNP positions, random genomic locations, and the non-replicate EMS-induced SNP positions, respectively. Of these, 38% (21,993 positions), 37% (20,156 positions) and 24% (14,099) of the error-prone, random genomic and the random EMS-induced SNPs, respectively, had perfect matching paralogous sequences in the reference genome. A similar distribution of imperfect matches was found for the three sets of sequences using a BLAST e-value threshold of e-10. Considering only end-to-end ungapped alignments of these 51bp contexts we detected a total of 68,568,482 close matches for 41,733 (73%) of the error-prone SNPs positions. For the randomly selected genomic positions, we detected ungapped near-perfect matches for nearly as many loci (33,980 corresponding to 62%) but in only 29,849,107 global alignments, indicating lower copy numbers for the paralogous sequences. This pattern of lower paralog coverage for the 51nt SNP context was also true for the non-replicate SNPs where we recover 3 fold fewer paralogous locations on average (21,870,737 global alignments for 33,605 sites corresponding to 58%). Interestingly, 51% (34,966,393 alignments) of the ungapped alignments to error-prone SNP sequence contexts contained a mismatch at the central SNP position. In contrast, only 7% (1,996,337 alignments) and 11% (2,369,537 alignments) of the ungapped alignments for the random genomic positions and the non-replicate EMS-induced SNPs, respectively, contained a mismatch at the central (SNP) position (Table 4). The total length of the super scaffolds that have not been assigned to any chromosome in sorghum genome reference assembly version 2.1, is 68Mb and accounts for 9.4% of the genome assembly. Yet, over 51% (17,735,536 alignments) of the global alignments with a central mismatch in the error-prone SNPs alignments data had super scaffold origins. For the non-replicate EMS-induced SNPs and random genomic positions datasets, only 32% (763,384 alignments) and 28% (563,042 alignments) respectively, of the global alignments with a central mismatch occurred in super scaffolds. The substitutions at the central mismatch position in the global alignments matched the SNP call for 58% of the error-prone SNP positions (33,083 SNPs) but only 34.5% of the non-replicate EMS-induced SNP positions (20,011 SNPs). The end-to-end ungapped (global) alignments may contain more than one mismatch position. Next we parsed the BLAST global alignment results to retain only alignments containing a single mismatch at the central (SNP) position. This identified 2,032,958 alignments for the error-prone SNP sequence contexts. Of these, 47% (951,217 alignments) were to super scaffold assemblies (Supplementary Data 20). We also obtained 113,515 and 120,547 global alignments for the random genome and EMS-induced SNPs datasets respectively but only 19% (22,011 alignments) and 28% (33,904 alignments) matched super scaffolds assemblies, respectively. This method was more stringent, single mismatch, and only recapitulated 26% (14,913 SNP positions) and 7% (4,191 SNP positions) of the error-prone and EMS-induced SNP positions, respectively (Figure 7). These results demonstrate that paralogous sequences from poorly assembled regions of the genome match the error prone sites detected as replicate SNPs in whole genome sequence data from the reference genotype. The DNA represented by these assemblies disproportionally contribute to the detection of false positives in shotgun sequencing experiments.

**Table4.**
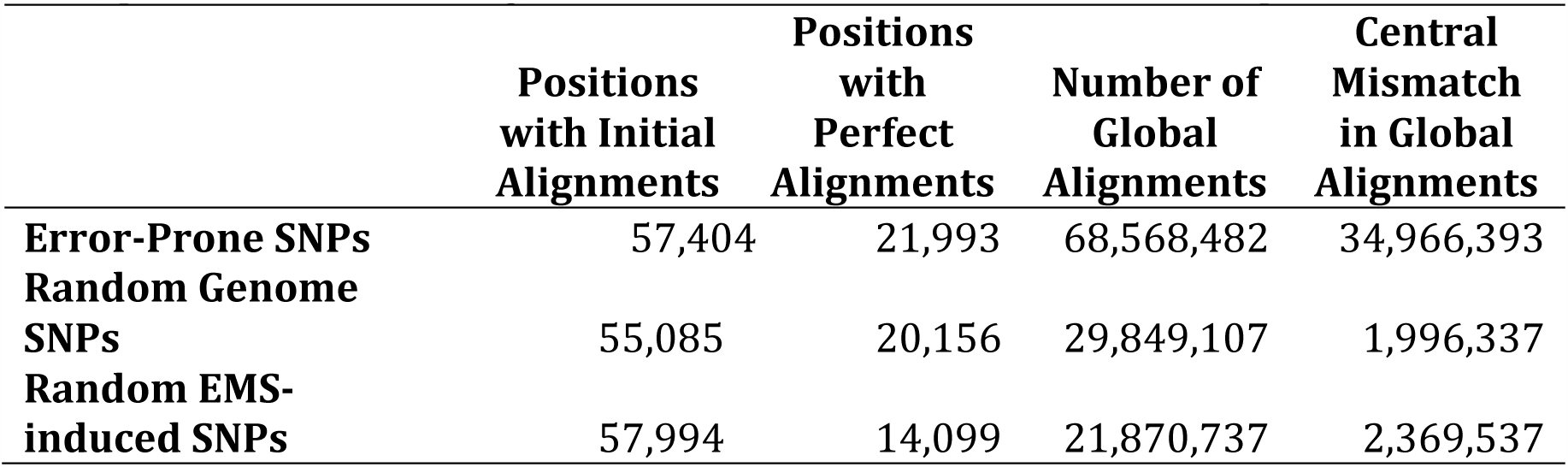
BLAST results for the global alignment of the 51nt sequence context of the error-prone SNPs, random genomic, and random EMS-induced SNPs positions.

**Figure 7.**
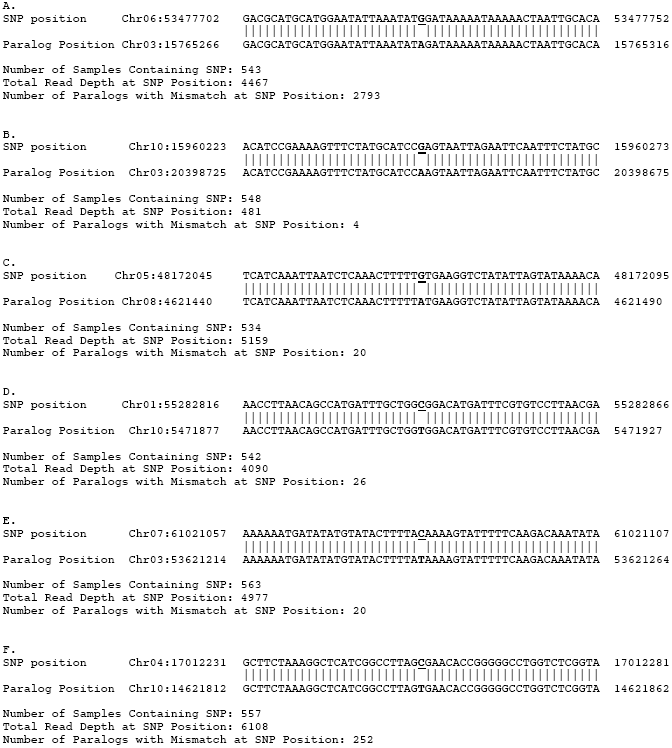
Examples of BLAST global alignments of the 51nt sequence context of the likely error-prone SNPs to the sorghum reference genome. The sample alignments depict paralogous alignments that result in false positive G:C to A:T EMS-induced mutations. The reference and SNP positions are highlighted in bold font, with the mutated position underlined.

This experimentally identifiable set of replicate SNP positions, which lack the expected EMS mutation spectrum, correspond to repeated sequences, contain paralogs with the predicted SNP and we refer to as the sorghum error prone positions (EPPs): the experimentally determined set of error prone positions. Similar approaches in any species should improve the accuracy of SNP calling by providing positions that can be screened out (Addo-Quaye et al., 2016).

### EMS-induced variants confirmation

A total of 29 randomly selected individuals from the collection of the 586 sequenced individuals were resequenced using leaf tissues from the M_5_ generation and the Illumina WideSeq paired-end sequencing protocol. The sequencing yielded mean genome coverage of 0.06X. These reads were aligned to the Sorghum genome and the positions of previously identified SNPs in these 29 lines were assessed. Given that a mean of 2,177 homozygous non-replicate SNPs were found in each line this should overlap with 130 SNPs per line and provide ample depth to test line identity. The low depth sequence overlapped with 3,843 non-replicate SNP positions and 96% of these were unambiguously confirmed (Supplementary Figure 9). This both validated our SNP calling procedures, and verified the identity of our lines and validity of our pedigrees. There were 1745 reads that contained positions called as heterozygous non-replicate SNPs in the M3 sequence data. For these heterozygous SNPs our low pass sequence confirmed 41% (710 out of 1,745) of the mutations (Supplementary Figure 10). This rate of validation again confirmed our pedigrees and line identities. The lower rate of confirmation is consistent with the expectations of segregation, where 37.5% of the M5 should now be homozygous reference for the heterozygous position and of the 25% still heterozygous, we will recover the non-reference allele in half of the reads. This should have resulted in 50% non-reference alleles or 48% if the validation rate was the same for heterozygous and homozygous SNPs. Consistent with the likely higher false positive rate of the heterozygous SNPs, suggested by the lower G:C to A:T SNP percentage in the non replicate SNPs, we validated fewer SNPs.

## DISCUSSION

The accuracy of the *in silico* detection of sequence variants can be impacted by the inherent sequencing error rates of NGS technologies, nature of the variant calling programs, genome complexities, sequence alignment programs and quality of reference genome assemblies (Cheng et al., 2014; Cheung et al., 2003; Estivill et al., 2002; Fredman et al., 2004; Nielsen et al., 2011; O’Rawe et al., 2013; Robasky et al., 2014). We propose a filtering step to improve current standard practices for filtering false positive variant calls detected in genetic studies using mutagenizing agents such as EMS, which induce random mutations. EMS mutagenesis should result in DNA mutations at nearly random subsets of the mutable positions in the genome of a treated organism. For an organism with a sufficiently large genome, the likelihood of a randomly induced mutation occurring at the same genomic position in two individuals that were treated independently is extremely low. If sequencing error, on the other hand, is non-randomly distributed, a SNP detected in multiple lines at the same genomic position is mostly likely a replicated false-positive SNP. On the other hand, a non-replicated high-quality SNP detected at a genomic position that is unique to a single individual in a population of independently EMS-mutagenized individuals is most likely a true positive and represents a real signal in the SNP detection. Applying this cost-effective and highly accurate filtering step to the large-scale NGS dataset produced from the M_3_ generation of an EMS-mutagenized population of the agronomically important crop species *Sorghum bicolor,* we showed that 76% of the homozgyous SNPs and 94% of the heterozygous SNPs detected using the standard SNPs detection and filtering were false positives (Figure 1 and Supplementary Figure 1). Thus, anyone applying this type of sequencing to identify a causal variant in the sequencing data from a single mutant line would have to sort through a majority of false positives. Two previous efforts had independently used a similar method to refine their variant calling (Sarin et al., 2010; Uchida, Sakamoto, Kurata, & Tasaka, 2011). However, our characterization and results clearly reveals empirical evidence of the accuracy and effectiveness of the approach. We identified and cataloged the relatively few genomic positions of the sorghum reference genome, consisting of 57,872 positions, that were most likely error-prone, and which produced 4,123,621 SNP calls of high statistical quality (Supplementary Data 2). Similar filtering of the insertion and deletion alleles retained less than 1% (17,760) as non-replicate indels (Supplementary Figure 3, Supplementary Table 3). Consistent with the analysis of indels induced by fast neutron irradiation of rice (G. Li et al., 2016), and the analysis method we employed, the vast majority of indels alleles were short in length (2.6 bases) with size ranging from 1 to 50 bases. In agreement with the high false positive detection rate for indels, the recent genome-wide analysis of chemically induced-mutations obtained by N-methyl-N’-nitro-N-nitrosoguanidine mutagenesis in the social soil amoeba *Dictyostelium discoideum* also uncovered very few likely true positive alleles, while discarding 99% of initially predicted variants (C.-L. F. Li, Santhanam, Webb, Zupan, & Shaulsky, 2016). Previous studies had revealed the potential for genome assembly errors in the presence of repetitive sequences and segmental duplications, that result in false positive or ambiguous variants (Cheung et al., 2003; Estivill et al., 2002; Fredman et al., 2004). Our examination of the sequence context of the identified error-prone positions showed evidence of paralogous sequence variants with origins in repetitive and duplicated genomic regions, contributed to false positive detections. Another source for the replicated SNP calls could be spontaneous mutations that arose in the parental line used in the mutagenesis but absent in the reference genome genotype. For example, a recent resequencing of the sorghum BTx623 parental line utilized in a mutagenesis experiment revealed 13,243 high confidence variations, when compared to the reference sorghum genotype (Jiao et al., 2016). We found that the average sequence coverage depths at the error-prone SNP positions and the bona fide SNP positions profiles were not different in our data. This may be due to the randomized assignment of non-unique reads from repeated sequences to each repeat location by the BWA aligner (Firtina & Alkan, 2016). Novel methods for improving the assignment of multi-mapping reads are emerging and may improve SNP calling when widely implemented (Johnson, Yeoh, Coruh, & Axtell, 2016).

The mutagenic effect of EMS is well-characterized in a wide variety of viruses, prokaryotes, fungal, plant, and animal genomes (Loveless, 1958; Loveless, 1969; Sega, 1984; Shrivastav et al., 2010; Singer, Kusmierek, & Ku, 1982). The predominant DNA reactive site that produces DNA lesions after EMS alkylation is O^6^ guanine. The ethylation of O^6^ guanine results in low mutagenicity in the EMS-treated organism (A. Loveless, 1969; Singer et al., 1982). We found an average of 2,177 homozygous and 815 heterozygous non-replicate SNP calls per sorghum individual (Table 3) and an estimated mutation rate of 4.0×10^−6^. The mispairing of O^6^ alkyl guanine with thymine results in G:C to A:T mutations (Kohalmi & Kunz, 1988; A. Loveless, 1969; Prakash & Sherman, 1973). The mutation spectra of the detected homozygous and heterozygous non-replicate SNPs showed 98% and 94% were G:C to A:T transitions, respectively. The slightly lower percentage of G:C to A:T transitions for the heterozygous SNPs may be attributed to higher false positive rates due to the inclusion of a greater number of sites with paralogous positions within the heterozygous SNP class as well as the lower overall stringency when non-SNP containing reads are allowed. The notion that we still have some false positives within our SNP calls is supported by the fact that within the protein-coding regions over 99% of the homozygous SNPs we detected were G:C to A:T substitutions. These positions are less likely to possess unassembled paralogous sequences than the intergenic positions. This frequency matches a previous study in Arabidopsis that found over 99% of EMS-induced mutations in a set of 192 genes were G:C to A:T substitutions (Greene et al., 2003). Although DNA lesions from EMS mutagenesis are predominantly associated with O^6^ guanine ethylation, other minor sites such as O^2^ and O^4^ thymine had also been suggested. Studies show the alkylation of O^4^ thymine results in mispairing with guanine and this leads to A:T to G:C transitions (Coulondre & Miller, 1977; Prakash & Sherman, 1973; Singer et al., 1978). Our analysis revealed that A:T to G:C transitions were the second most abundant class of likely EMS-induced mutations and accounted for 0.7% of the homozygous SNPs. This observation corroborates the finding in viruses that suggests EMS produces A:T to G:C transitions at about two orders of magnitude less frequently than G:C to A:T transitions (Krieg, 1963).

We examined the sequence context of the EMS-induced SNPs, for possible evidence of conserved motifs flanking the mutated genomic positions. If detected, this indicates a non-randomness of EMS mutagenesis beyond the chemical preference for alkylating guanine. Previous large-scale analysis of flanking positions of predicted EMS-induced mutations in sorghum reported an overrepresentation of cytosine residues at positions −2 and +1 (Jiao et al., 2016). In addition to these sites, we detected a more extensive signal characterized by a genome-wide overrepresentation of a 5’-CGCG-3’ motif and an underrepresentation of a 5’-AHGM-3’ motif (Figure 6 and Supplementary Figures 7 and 8). The CG dinucleotide content in the 21nt context of the mutation was 57% higher than that of the sorghum reference genome. The observation that a similar sequence occurs as an overrepresented motif was described previously in the large-scale exome sequencing of rice EMS mutants (Henry et al., 2014). This observation suggests that something in common between the two systems, either inherent to the structure of that DNA sequence, the repair process, or the methylation of cytosines is responsible for the higher rate of mutation. Whereas the rice EMS-induced mutations were more likely at unmethylated cytosines (Henry et al., 2014), sorghum EMS-induced mutations were overrepresented at methylated cytosines (Jiao et al., 2016). It is important to note, that while the relationships between cytosine methylation and EMS-induced mutations were in opposite directions in rice and sorghum, methylation did not affect a majority of positions.

A total of 4,220 transcripts corresponding to 3,736 protein-coding genes lacked any predicted mutations. Given the total number of mutations we did not expect to return mutations in all genes. As expected, the short genes with few G:C sites were more likely to have no mutations in our population. Interestingly, in sorghum we found no direct correlation between GC-content and the number of SNPs in a transcript (Figure 5A). However we observed linear correlations between the number of SNPs in a transcript and the G+C-count and also with transcript length (Figure 5B and 5C). In effect, since EMS mutagenesis targets predominantly random G positions in the genome, the number of mutations in a gene is correlated with the gene length and the G+C-count. Due to the nucleotide composition of the three canonical translational stop codons (TAA, TGA and TAG), and the fact that the majority of the likely EMS-induced mutations are G:C to A:T substitutions, stop codons are primarily buffered from the effects of EMS mutagenesis and we saw very few losses of stop codons. TAA codons do not contain any G and/or C nucleotide targets of EMS, while the G to A substitution of TGA or TAG converts them silently to TAA stop codons. We did observe 148 such silent stop codon substitutions in our SNPs annotation (Supplementary Table 6), which may be of use to others interested in the regulation of stop codons, the contribution of stop codon usage to mRNA turnover, or nonsense suppression. We observed a single homozygous and four heterozygous stop codon loss SNPs which were due to non-G:C to A:T substitutions that should result in translation read-through (Table 3).

In biological experiments, replication plays a critical role by adding statistical power to both detect differences and reject errant stochastic results, as discussed previously for NGS experiments (Robasky et al., 2014). We leveraged replication to detect, among standard filtered DNA variants in independently mutagenized lines, signals of false positive SNP calls. This improved the accuracy of our predictions by many-fold. In the absence of inter-sample variant comparisons, the majority of the standard filtered SNPs in each line will appear to be possible causative mutations. If the same line were to be resequenced, it would return a subset of the false positive calls a second time, meeting the requirement of reproducibility, and these will be falsely confirmed as true DNA variants. This problem points to a need within all research communities to share primary sequence data and SNP calls more openly to improve forward genetics and gene functional characterization. Each lab that is using next generation sequencing to map mutants is sequencing lines of independent pedigrees. Without biological replication, each lab in isolation is calling these reproducibly errant SNP calls and including them in the set of possible causative SNPs in their experiment. By sharing SNP calls, and raw sequence data, from all experiments sequencing mutants underway in a scientific community, replication of sequencing can be achieved without additional cost to each lab and funding agency. By identifying the errant-prone positions collaboratively, all labs no longer need to follow up on SNPs that cannot be responsible for the mutation they seek to describe. An open data culture would permit current research efforts to more efficiently discover alleles responsible for mutant phenotypes in any genetic model system.

Although the removal of the shared variants is a very effective technique for identifying likely false positive EMS-induced mutations, a few caveats need to be highlighted. Firstly, the method should be applied meticulously to prevent the introduction of false negatives. For example, sample mislabeling and unexpected hybridization between individuals within the population can result in the sequencing of multiple DNA samples containing shared (identity by descent) mutagenized chromosomes. Also sequence contamination originating from sample or library preparation stages can negatively impact the detection of false positives. In all the above-mentioned scenarios, technical errors or contaminants would result in the erroneous purging of true DNA variants that were shared due to identity by descent from a common ancestor. These errors are detectable. In contrast to non-contaminated samples, prior to population-wide subtraction of shared variations, sample pairs connected by contamination are characterized by an abnormally high percentage of shared variation between the two individuals. Prior to the removal of common variants, pairwise comparison of detected variants in all the sequenced individuals in the population is mandated to expose contaminated samples with shared pedigrees, regardless of how certain biologists are that there is a lack of contamination in their work. Secondly, although the likelihood is extremely low, it is not statistically impossible for a mutation to be independently induced at the same genomic position in two or more individuals. Hence it is noteworthy that there is a possibility that a few replicate variants are real. This frequency increases as the sample size and mutation density go up and as mutations are non-randomly distributed in the genome.

The finding that the replicate SNPs were present in poorly assembled regions of the genome and present in a greater number of paralogs has a number of implications (Figure 7). Firstly, the poor assembly that results in unplaced scaffolds differentially affects repeated sequences. Much of the SNP calling may actually result from the sequence derived from DNA not correctly represented in the sorghum genome assembly which is only 727 Mb while the genome is 830 Mb (Arumuganathan & Earle, 1991; Peterson et al., 2002; Price et al., 2005). These unknown sequences are extracted and sequenced with every experiment but their appropriate mapping is impossible to know. We suspect that they contribute significantly to the error prone positions and that genomes with less unassembled and less poorly scaffolded DNA will produce fewer false positive SNPs. We also predict that a consequence of presence-absence variation between individuals will be the production of lineage-specific error prone SNPs. The locations of the presence/absence (PAV) should be determinable by mapping the SNPs produced by misalignment of PAV-derived DNA. Another problem in genome structure is indicated by the difference in paralog detection for randomly selected positions versus our non-replicate SNP positions (Table 4). The non-replicate SNP positions were drawn from a dramatically lower proportion of positions with perfect match paralogs in the genome than expected by random chance (24% in the non-replicate SNPs but 37% of randomly selected genomic locations). This indicates a strong negative effect of a perfect match paralog on homozygous SNP detection, the false negative flip-side of paralogy to the error prone positions. This demonstrates that SNP calling in polyploids, even paleopolyloids, will be suppressed at positions of conservation by ambiguous alignment and the erroneous assessment of no variation. Special procedures for calling SNPs at paralogs should improve SNP detection in subsets of genomes characterized by identity. The quality of reference genome assemblies, the complexity of genomes, and a careful, critical view of next generation sequence data demonstrate the need for a more sophisticated approach to false positive detection when rare variant identification is the experimental goal. Based on our work in a homozygous model system, we are deeply skeptical of rare variant detection claims that have not sought trivial explanations for the repeated detection of SNPs in experimental samples. For instance, the relevance of the error prone positions to the cataloging of rare variants induced during cancer progression is needed.

## CONCLUSION

The prevalence of false positive DNA variants negatively impacts the efficiency of forward and reverse genetics studies. We proposed and described a cost-effective, high throughput and a highly accurate NGS method for improving the *in silico* detection of induced mutations via the identification of false positive variants. The proposed method is generally applicable to any genetics study, and is also independent of the choice of the variant calling program. Using the method, we have detected and described the set of EMS-induced mutations in a population of sorghum bicolor, which is a next-generation crop species with tremendous agronomical potential. We hope the genetics community and plant breeders will find this proposed method and the described genetic resource to be very useful.

## Supplementary data files

The supplementary data files for this project are available from the Dryad Digital Repository:http://datadryad.org/review?doi=doi:10.5061/dryad.t80gj

## Accession numbers

The Illumina next generation sequencing datasets generated for this project have been deposited in the NCBI SRA (Short Reads Archive), under the Project Title: *“Purdue University: Functional Gene Function Discovery Platform for Sorghum”*. The SRA Study Accession Number SRP065118. The BioProject ID# is PRJNA297450 and the BioSample ID#s range from SAMN04128840 to SAMN04129425. Seed samples for the EMS-mutagenized sorghum population used in this study are available on the Germplasm Resources Information Network (GRIN) database.

## Author contributions

MT, CW, CAQ and BPD designed the experiments.

CAQ and BPD performed the bioinformatics analysis.

CAQ and NC confirmed the detected SNPs CAQ and BPD drafted the manuscript.

BPD, MT, CW, CAQ and NC edited the manuscript.

## Acknowledgements

We thank Damon Lisch for helpful comments on this manuscript. We also thank Phillip SanMiguel and Rick Westerman of the Purdue University Genomics Core Facility for the NGS sequencing and data warehousing support. Finally, we also thank the IT staff of the Purdue University Rosen Center for Advanced Computing (RCAC) for the research computing and cyber infrastructure support.

## List of Supplementary Data Files

1. Supplementary Data 1: Standard filtered SNPs detected in each of the 586 sorghum individuals.

2. Supplementary Data 2: Probable error-prone SNP genomic positions in the sorghum reference genome (version 2.1).

3. Supplementary Data 3: Non-replicate, and likely EMS-induced homozygous SNPs in each of the 586 sorghum individuals.

4. Supplementary Data 4: Non-replicate, and likely EMS-induced heterozygous SNPs in each of the 586 sorghum individuals.

5. Supplementary Data 5: Counts of homozygous and heterozygous SNPs in all the 586 sequenced lines.

6. Supplementary Data 6: SnpEff functional classification of the homozygous EMS-induced SNPs in all the 586 sorghum individuals.

7. Supplementary Data 7: Function description for the genes containing the SnpEff-annotated homozygous EMS-induced SNPs in all the 586 sorghum individuals.

8. Supplementary Data 8: List of SnpEff-annotated EMS-induced SNPs predicted to trigger silent stop codon lost substitutions in genes of the 586 sorghum individuals.

9. Supplementary Data 9: SnpEff functional classification of the heterozygous EMS-induced SNPs in all the 586 sorghum individuals.

10. Supplementary Data 10: Function description for the genes containing the SnpEff-annotated heterozygous EMS-induced SNPs in all the 586 sorghum individuals.

11. Supplementary Data 11: SnpEff functional classification of the likely EMS-induced indels in all the 586 sorghum individuals.

12. Supplementary Data 12: Function description for the genes containing the SnpEff-annotated likely EMS-induced indels in all the 586 sorghum individuals.

13. Supplementary Data 13: SIFT prediction results for the missense-annotated homozygous EMS-induced SNPs in all the 586 sequenced individuals.

14. Supplementary Data 14: SIFT prediction results for the missense-annotated heterozygous EMS-induced SNPs in all the 586 sequenced individuals.

15. Supplementary Data 15: Summary statistics for the sequencing, mapping, variants prediction, filtering, annotation and classification for the detected EMS-induced variants in all 586 sequenced individuals.

16. Supplementary Data 16: Detailed classification, gene function description and annotation of the medium or high impact homozygous EMS-induced SNPs.

17. Supplementary Data 17: Detailed classification, gene function description and annotation of the medium or high impact heterozygous EMS-induced SNPs.

18. Supplementary Data 18: Detailed classification, gene function description and annotation of the medium or high impact EMS-induced indels.

19. Supplementary Data 19: The detailed attributes of gene transcripts, including number of SNPs, GC-content, sequence length, mononucleotide and dinucleotide counts, which were used in the linear correlation analysis.

20. Supplementary Data 20: The subset of BLAST ungapped global alignments for the 51nt sequence contexts of the error-prone SNP positions, randomly selected EMS-induced SNP positions and randomly selected genome positions.

21. Supplementary Data 21: Sample scripts for the variants detection and annotation pipeline.

**Supplementary Figure 1.**
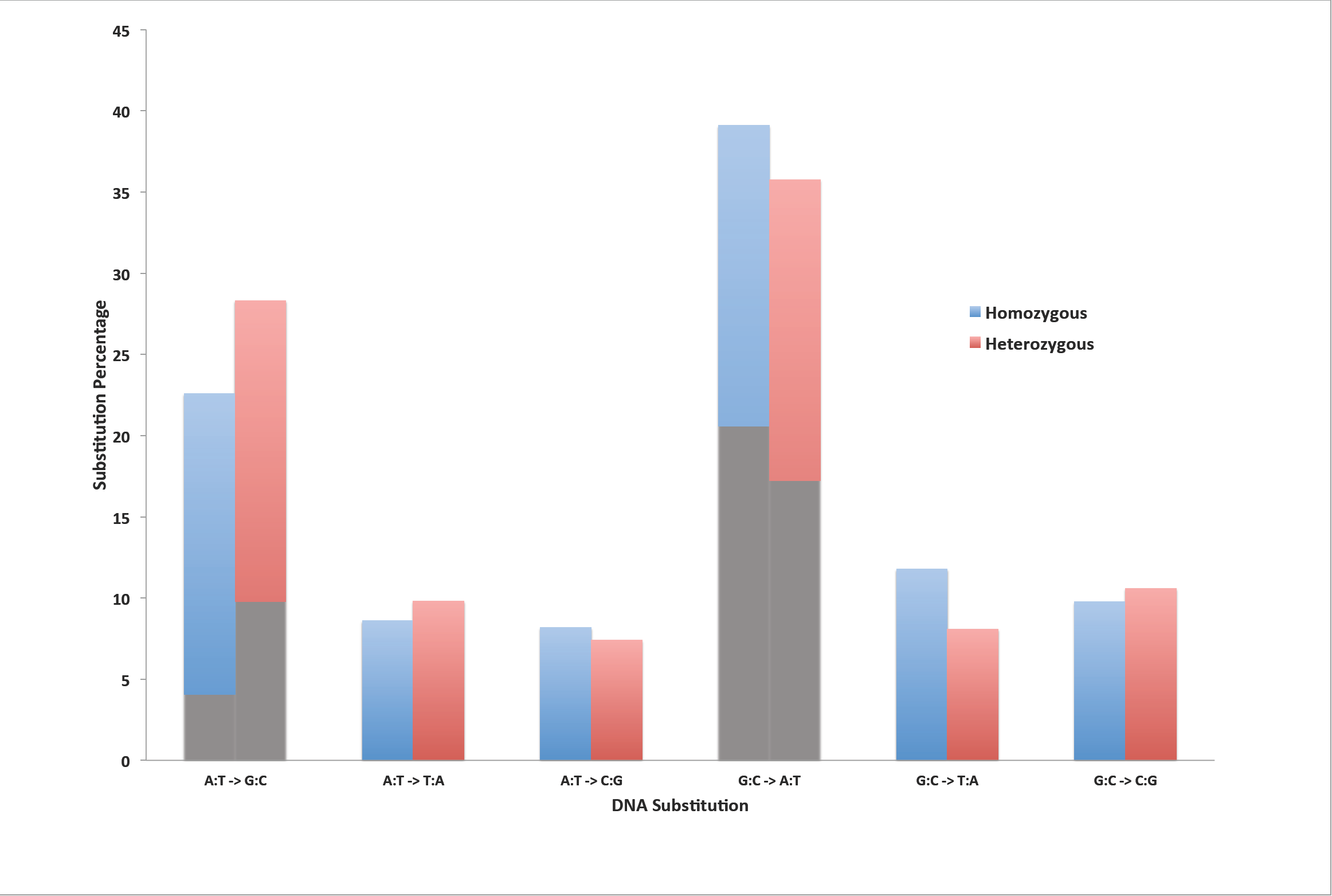
The mutation spectrum of the standard filtered SNPs predicted for the sorghum EMS-mutagenized population. The majority (61% of the homozygous and 64% of the heterozygous) of the SNPs were not G:C to A:T transitions and hence not likely to be products of EMS mutagenesis.

**Supplementary Figure 2.**
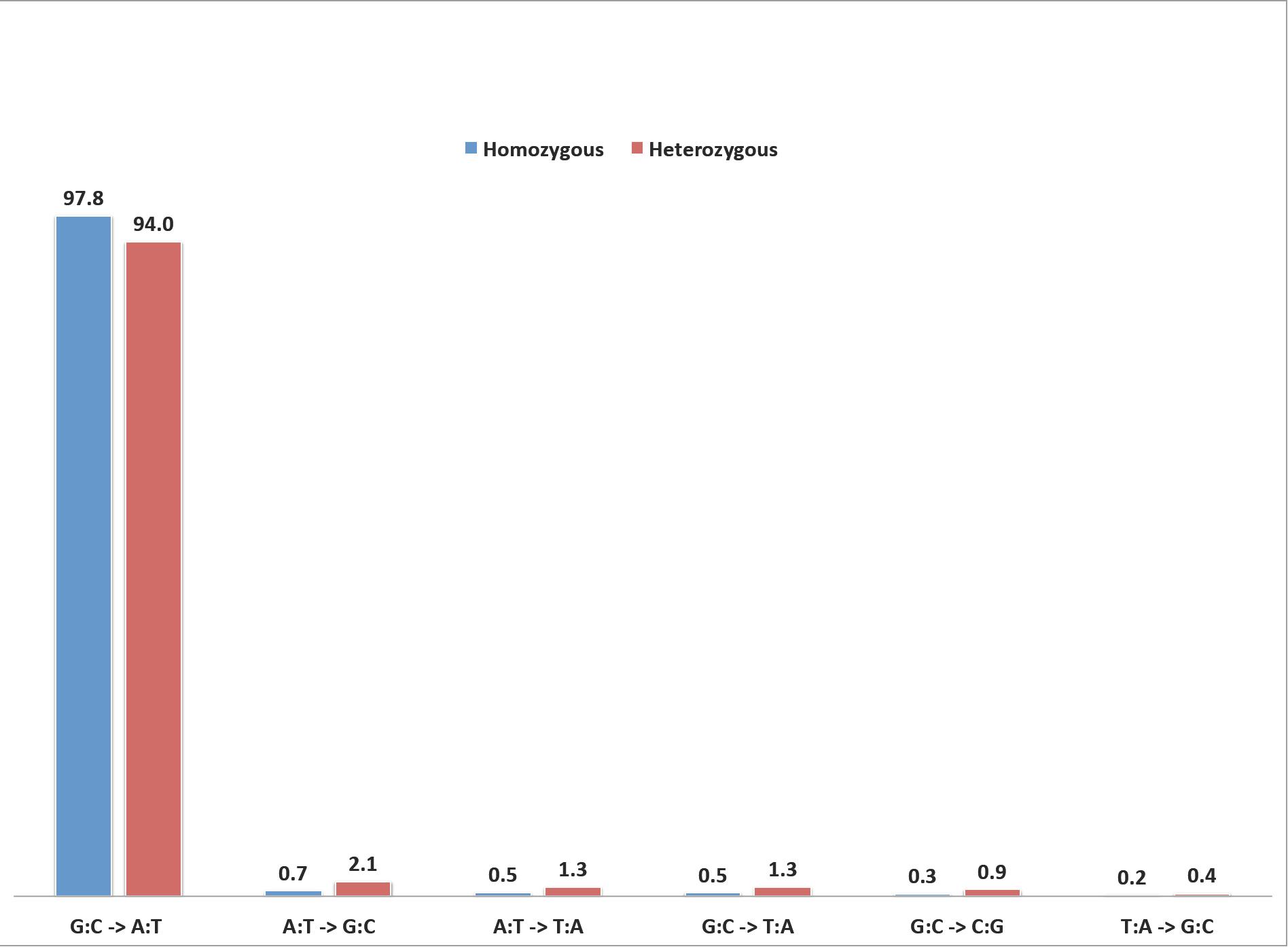
The mutation spectrum of the non-replicate homozygous and heterozygous SNPs detected in the 570 EMS-treated sorghum individuals. Removal of the replicate SNPs with origin in the error-prone genomic positions resulted in predominantly G:C to A:T substitutions. The non-replicate SNPs are most likely EMS-induced SNPs.

**Supplementary Figure 3.**
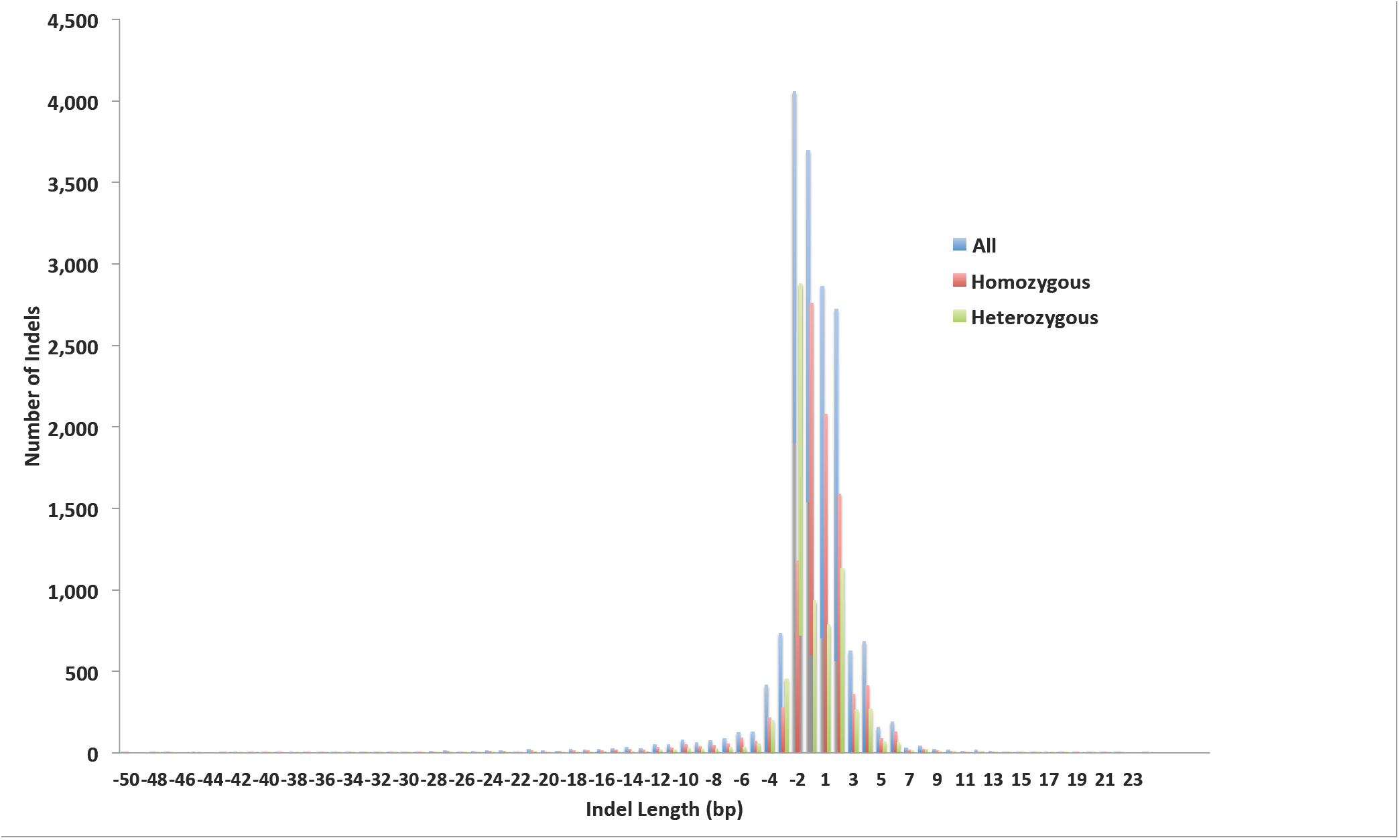
The length distribution of the predicted EMS-induced insertion/deletion polymorphisms in the 586 EMS-mutagenized sorghum individuals. The deletions are depicted as negative lengths and insertions represented by positive lengths.

**Supplementary Figure 4.**
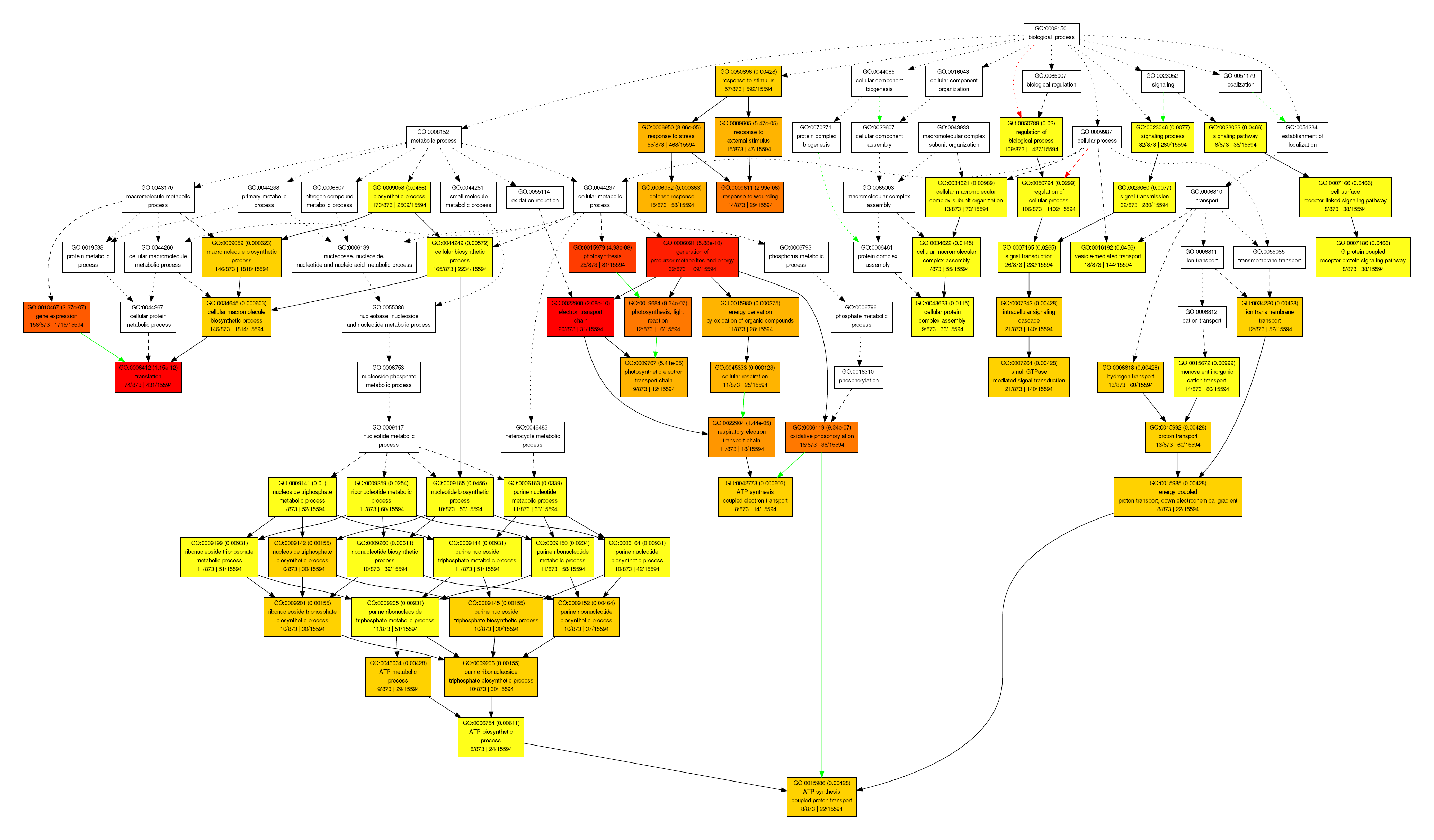
Gene ontology annotation of the predicted biological functions of the subset of sorghum genes with no predicted EMS-induced mutations in the resequenced 586 EMS-mutagenized individuals.

**Supplementary Figure 5.**
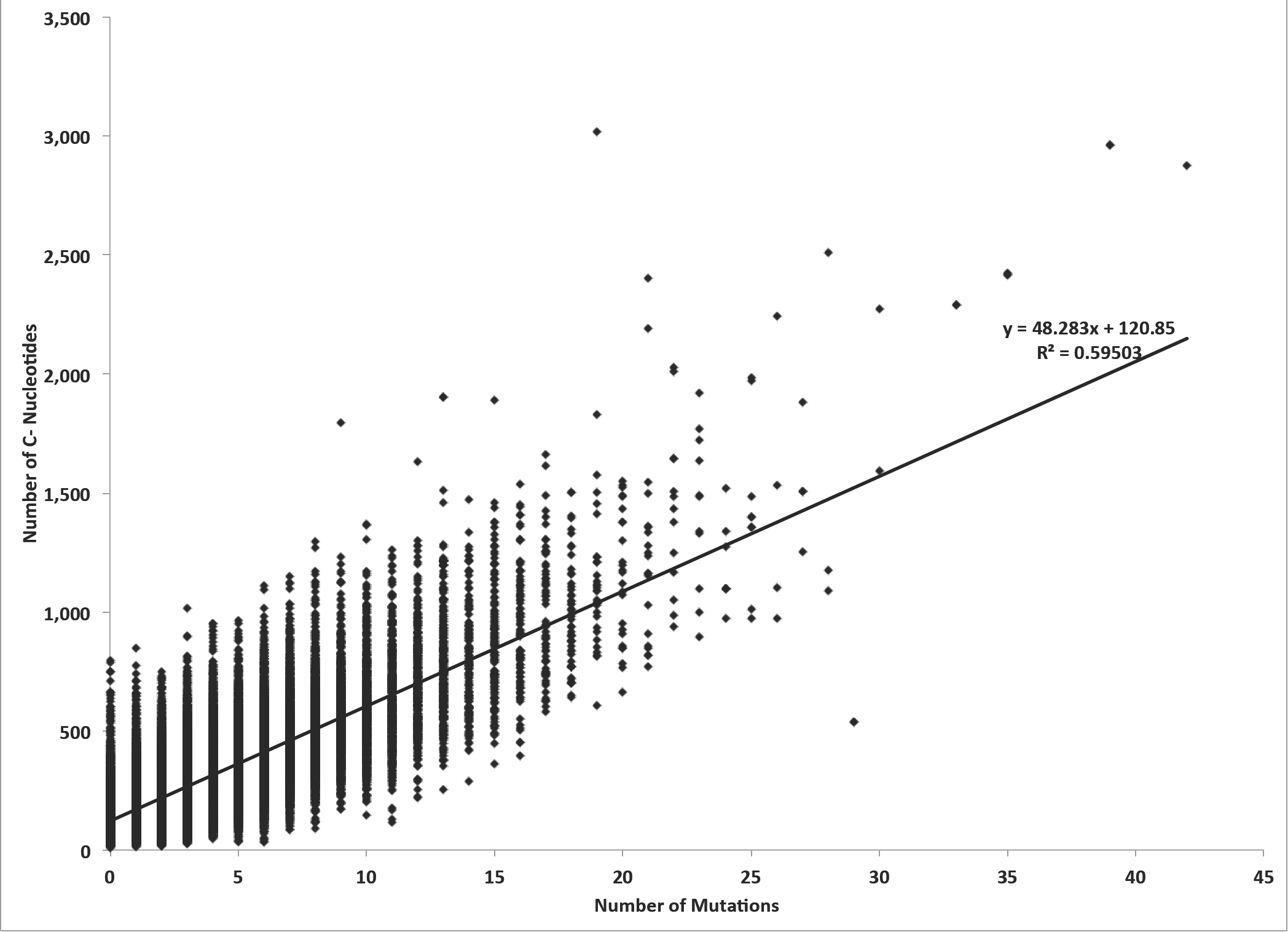
The linear correlation between the predicted number of EMS-induced mutations and the total number of G-nucleotide residues in a gene transcript.

**Supplementary Figure 6.**
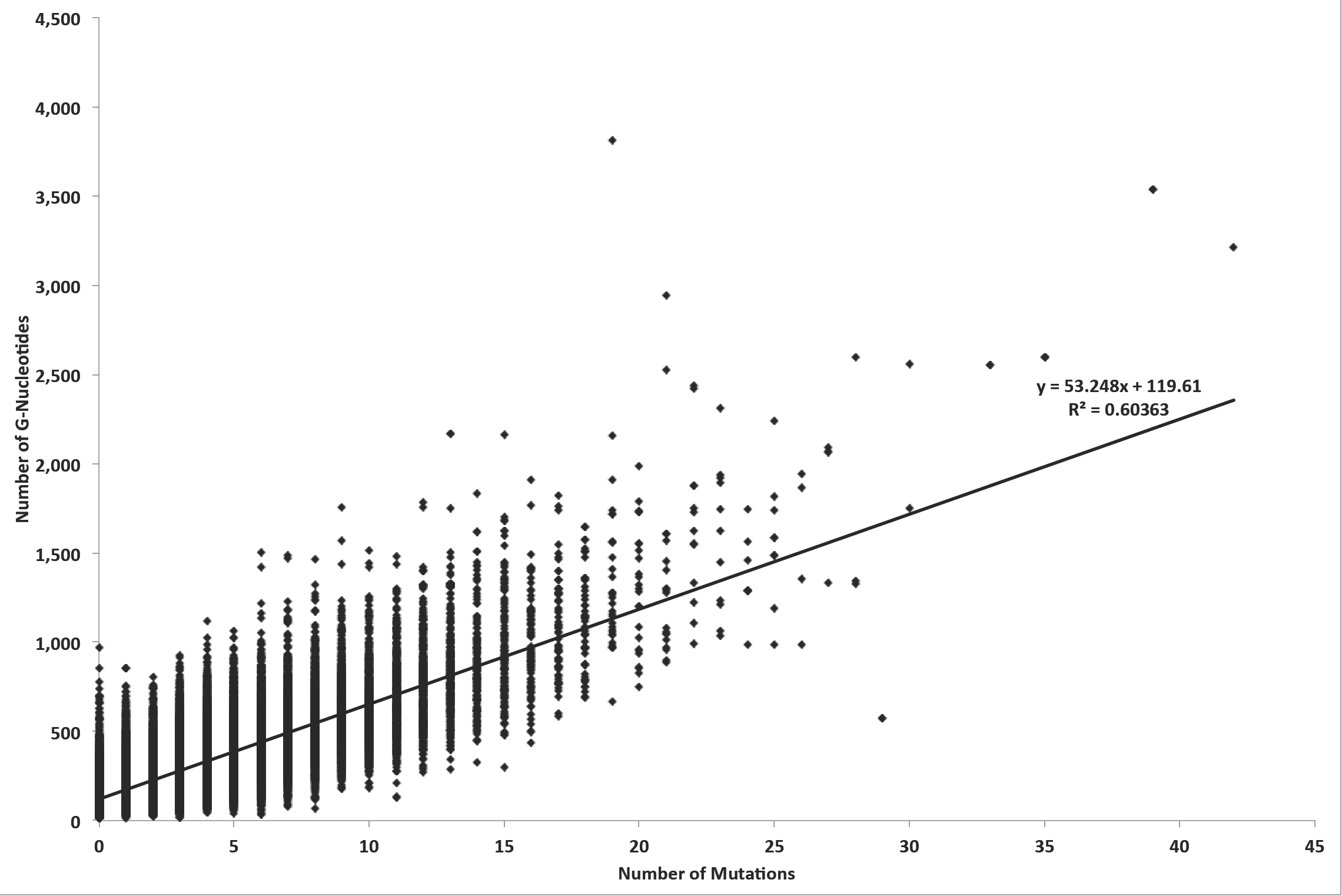
The linear correlation between the predicted number of EMS-induced mutations and the total number of C-nucleotide residues in a gene transcript.

**Supplementary Figure 7.**
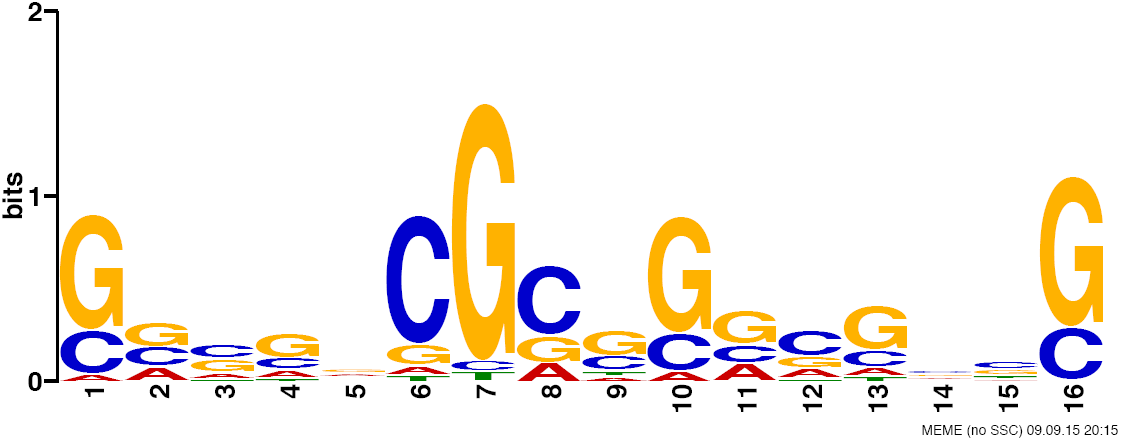
Analysis of the 21nt sequence context of 10,000 randomly selected EMS-induced homozygous mutations in the 586 resequenced *sorghum bicolor* genomes. Using the MEME unngapped motif finder the topmost motif (p-value of 5.7e-304) is 16 bases long and is predominantly CG-rich. The predominant EMS-induced mutational target is the G-nucleotide at position 7.

**Supplementary Figure 8.**
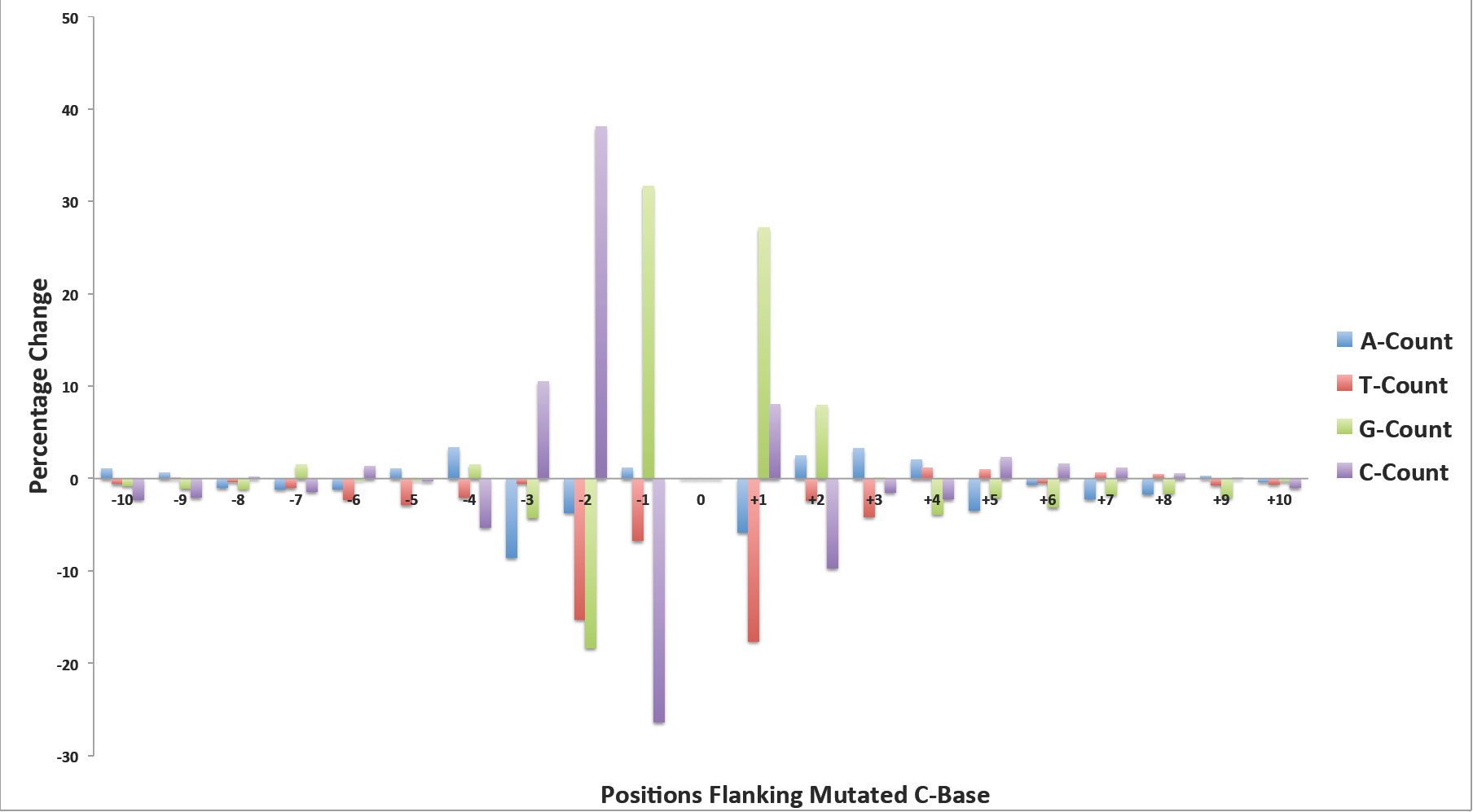
The 21nt sequence contexts comparison for the mutated C-residue DNA sites in the EMS-induced SNPs and randomly selected positions in the sorghum reference genome. The percentage change in the nucleotide frequency for all DNA base types in all DNA positions shows an overrepresentation of a 5’ -CG**C**G-3’ motif. The mutated C-residue is shown in bold.

**Supplementary Figure 9.**
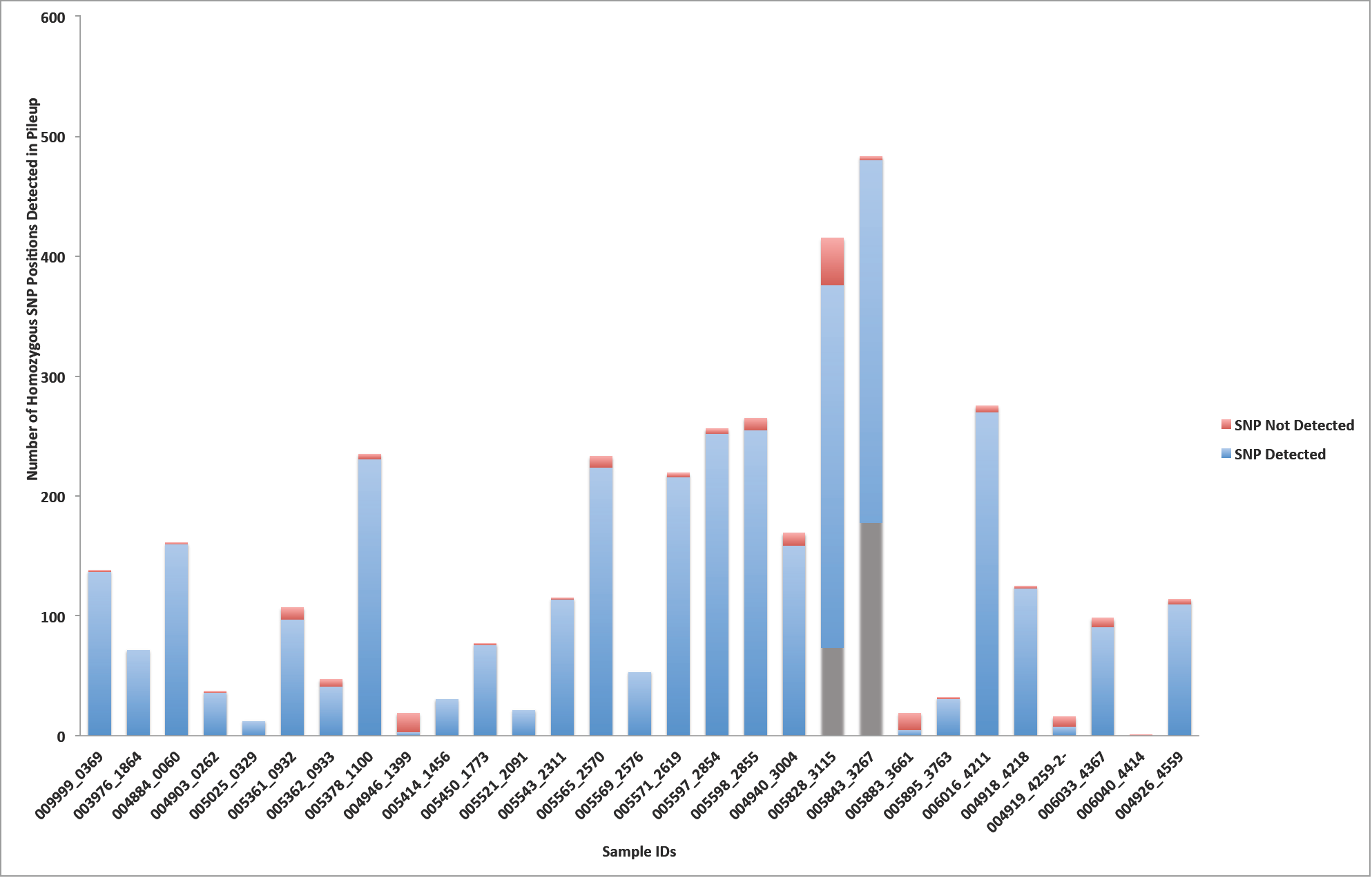
Confirmation of the EMS-induced homozygous SNP positions in the resequenced genomes of the M_5_ generation of 29 randomly selected subset of the original 586 EMS-mutagenized sorghum individuals.

**Supplementary Figure 10.**
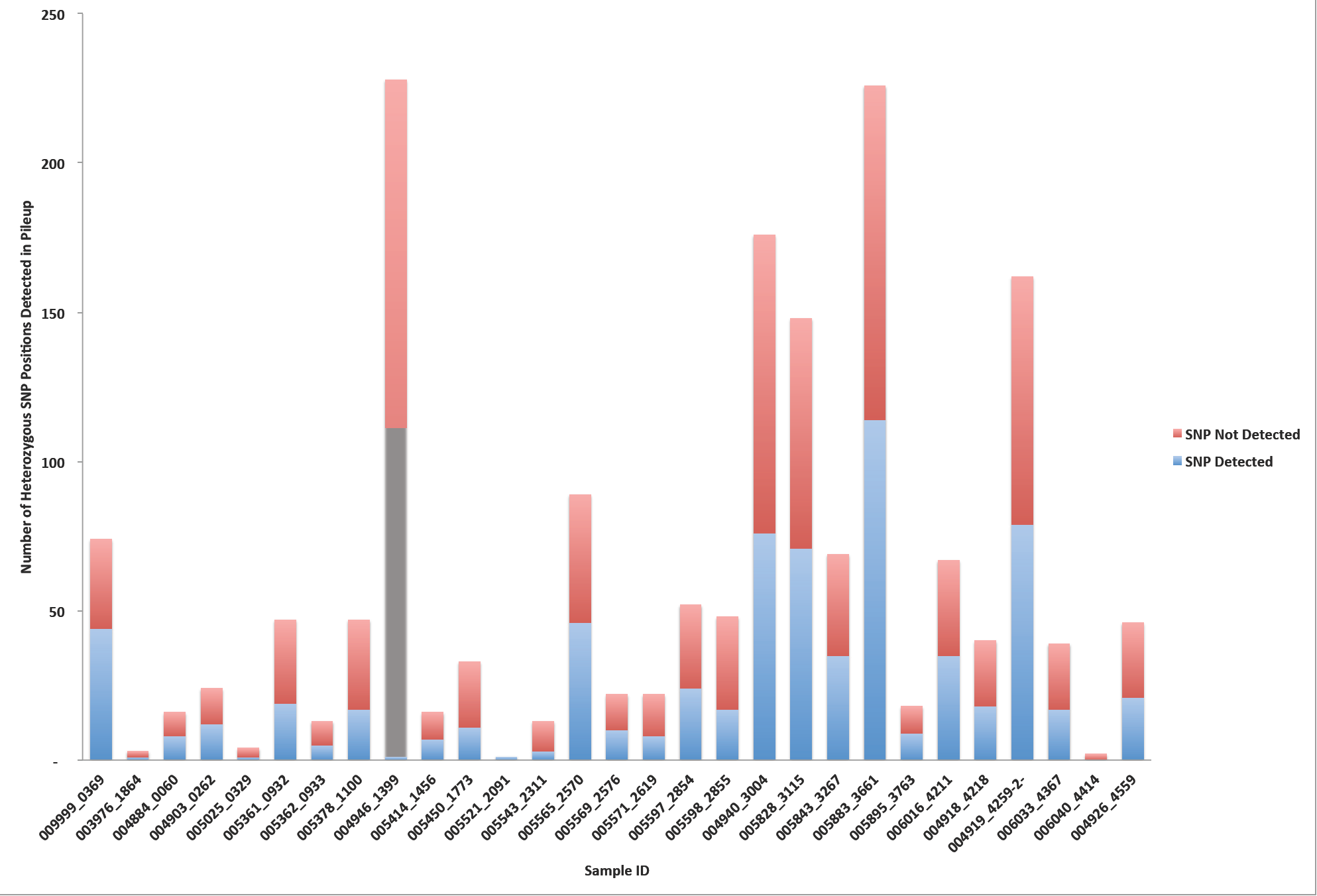
Confirmation of the EMS-induced heterozygous SNP positions in the resequenced genomes of the M5 generation of 29 randomly selected subset of the original 586 EMS-mutagenized sorghum individuals.

**Supplementary Table 1.**
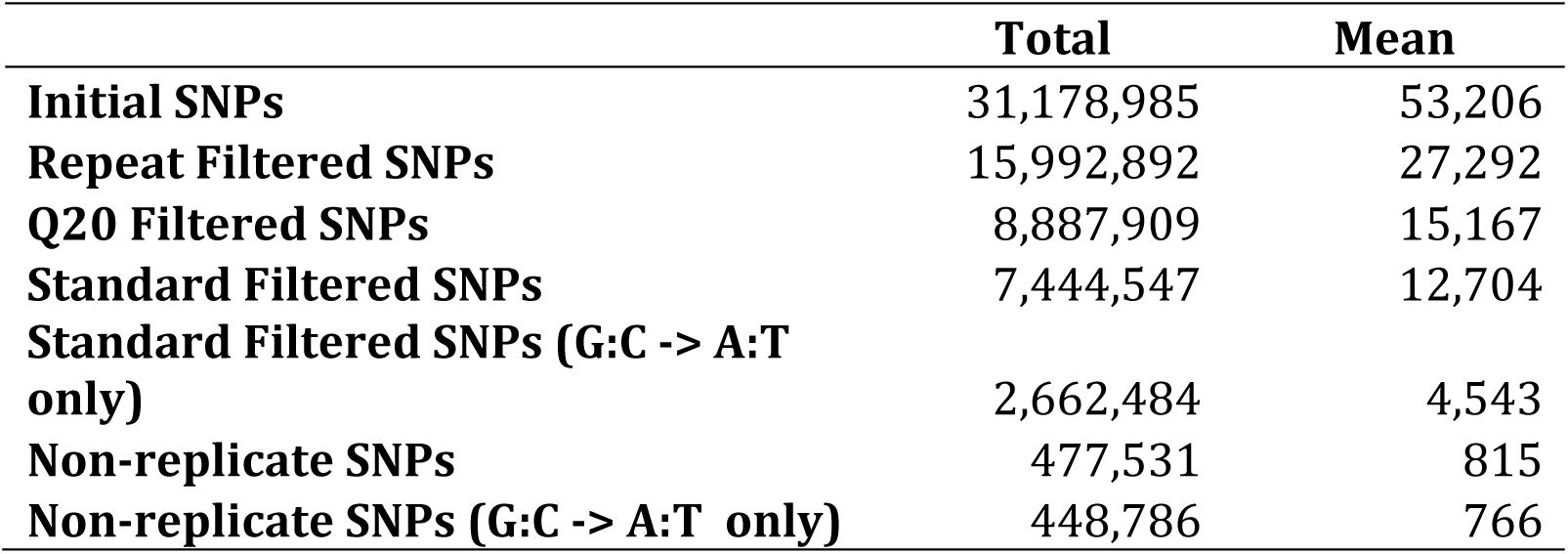
Summary statistics for the detection of heterozygous single nucleotide polymorphisms in the 586 EMS-mutagenized sorghum BTx623 individuals.

|  | **Total** | **Mean** |
| --- | --- | --- |
| **Initial SNPs** | 31,178,985 | 53,206 |
| **Repeat Filtered SNPs** | 15,992,892 | 27,292 |
| **Q20 Filtered SNPs** | 8,887,909 | 15,167 |
| **Standard Filtered SNPs** | 7,444,547 | 12,704 |
| **Standard Filtered SNPs (G:C -> A:T**<br>**only)** | 2,662,484 | 4,543 |
| **Non-replicate SNPs** | 477,531 | 815 |
| **Non-replicate SNPs (G:C -> A:T only)** | 448,786 | 766 |

**Supplementary Table 2.** Frequency distribution of genomic positions for replicate SNPs predicted repeatedly in the 570 EMS-treated sorghum individuals.

| **Number of Individuals** | **Number of**<br>**Genomic**<br>**Positions** | **SNPs**<br>**Count** |
| --- | --- | --- |
| 2-50 | 42,424 | 356,850 |
| 51-100 | 3,545 | 256,951 |
| 101-150 | 2,283 | 284,416 |
| 151-200 | 1,712 | 298,498 |
| 201-250 | 1,466 | 329,905 |
| 251-300 | 1,285 | 352,453 |
| 301-350 | 1,231 | 400,990 |
| 351-400 | 1,126 | 422,004 |
| 401-450 | 1,037 | 440,956 |
| 451-500 | 1,085 | 515,902 |
| 501-570 | 678 | 352,488 |
| **Total** | **57,872** | **4,011,413** |

**Supplementary Table 3.** Mutation spectrum of the standard filtered SNP calls in The 570EMS-mutagenized sorghum BTx623 individuals.

|  | Homozygous SNPs |  |  | Heterozygous SNPs |  |  |
| --- | --- | --- | --- | --- | --- | --- |
|  | All | Redundant | Unique | All | Redundant | Unique |
| A->G | 589,986 | 585,456 | 4,530 | 1,027,490 | 1,022,616 | 4,874 |
| A->T | 222,831 | 219,766 | 3,065 | 340,402 | 337,464 | 2,938 |
| A->C | 208,555 | 207,313 | 1,242 | 269,942 | 268,967 | 975 |
| G->A | 1,027,845 | 424,322 | 603,523 | 1,272,951 | 1,057,127 | 215,824 |
| G->T | 289,557 | 286,635 | 2,922 | 287,019 | 284,052 | 2,967 |
| G->C | 268,467 | 266,427 | 2,040 | 386,398 | 384,396 | 2,002 |
| T->A | 228,101 | 224,972 | 3,129 | 370,442 | 367,375 | 3,067 |
| T->G | 220,384 | 219,170 | 1,214 | 266,145 | 265,107 | 1,038 |
| T->C | 594,495 | 589,959 | 4,536 | 1,022,416 | 1,017,636 | 4,780 |
| C->A | 328,243 | 325,332 | 2,911 | 298,493 | 295,461 | 3,032 |
| C->G | 243,391 | 241,357 | 2,034 | 380,364 | 378,290 | 2,074 |
| C->T | 1,022,321 | 420,704 | 601,617 | 1,315,553 | 1,095,963 | 219,590 |
| **Total** | **5,244,176** | **4,011,413** | **1,232,763** | **7,237,615** | **6,774,454** | **463,161** |

**Supplementary Table 4.** Mutation spectrum of the standard filtered SNP calls in the 586 EMS-mutagenized sorghum BTx623 individuals.

|  | Homozygous SNPs |  |  | Heterozygous SNPs |  |  |
| --- | --- | --- | --- | --- | --- | --- |
|  | All | Redundant | Unique | All | Redundant | Unique |
| A->G | 606,561 | 601,853 | 4,708 | 1,056,494 | 1,051,453 | 5,041 |
| A->T | 229,188 | 226,017 | 3,171 | 350,218 | 347,181 | 3,037 |
| A->C | 214,225 | 212,933 | 1,292 | 277,673 | 276,661 | 1,012 |
| G->A | 1,061,030 | 436,074 | 624,956 | 1,309,550 | 1,087,015 | 222,535 |
| G->T | 297,774 | 294,735 | 3,039 | 295,284 | 292,237 | 3,047 |
| G->C | 275,947 | 273,827 | 2,120 | 397,443 | 395,355 | 2,088 |
| T->A | 234,469 | 231,219 | 3,250 | 380,959 | 377,790 | 3,169 |
| T->G | 226,608 | 225,343 | 1,265 | 273,641 | 272,565 | 1,076 |
| T->C | 611,321 | 606,628 | 4,693 | 1,051,284 | 1,046,318 | 4,966 |
| C->A | 337,551 | 334,526 | 3,025 | 306,979 | 303,844 | 3,135 |
| C->G | 250,213 | 248,091 | 2,122 | 391,298 | 389,141 | 2,157 |
| C->T | 1,054,606 | 432,375 | 622,231 | 1,352,934 | 1,126,683 | 226,251 |
| **Total** | **5,399,493** | **4,123,621** | **1,275,872** | **7,443,757** | **6,966,243** | **477,514** |

**Supplementary Table 5.** Classification of the predicted EMS-induced SNPs and their effects in the 570 EMS-mutagenized sorghum BTx623 individuals

| SNPs Classification | Total |  | Mean |  |
| --- | --- | --- | --- | --- |
|  | Homozygous | Heterozygous | Homozygous | Heterozygous |
| Stop Gained | 2,776 | 1,317 | 5 | 2 |
| Splice Site Donor | 588 | 298 | 1 | 1 |
| Splice Site Acceptor | 413 | 101 | 1 | 0 |
| Start Lost | 107 | 33 | 0 | 0 |
| Stop Lost | 1 | 3 | 0 | 0 |
| High Impact | 3,885 | 1,752 | 7 | 3 |
| Moderate Impact | 54,412 | 19,277 | 95 | 34 |
| Start Gained | 3,255 | 1,202 | 6 | 2 |
| 5' Prime UTR | 15,714 | 5,212 | 28 | 9 |
| 3' Prime UTR | 24,176 | 9,581 | 42 | 17 |
| Introns | 105,307 | 41,118 | 185 | 72 |
| Intergenic | 997,889 | 375,723 | 1,751 | 659 |

**Supplementary Table 6.**
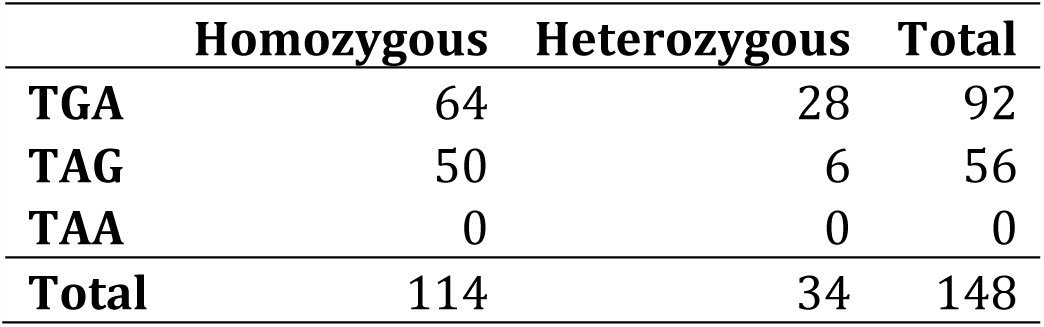
Distribution of synonymous stop codon lost substitutions in the 586 EMS-mutagenized sorghum BTx623 individuals.

|  | Homozygous | Heterozygous | Total |
| --- | --- | --- | --- |
| TGA | 64 | 28 | 92 |
| TAG | 50 | 6 | 56 |
| TAA | 0 | 0 | 0 |
| Total | 114 | 34 | 148 |

**Supplementary Table 7.**
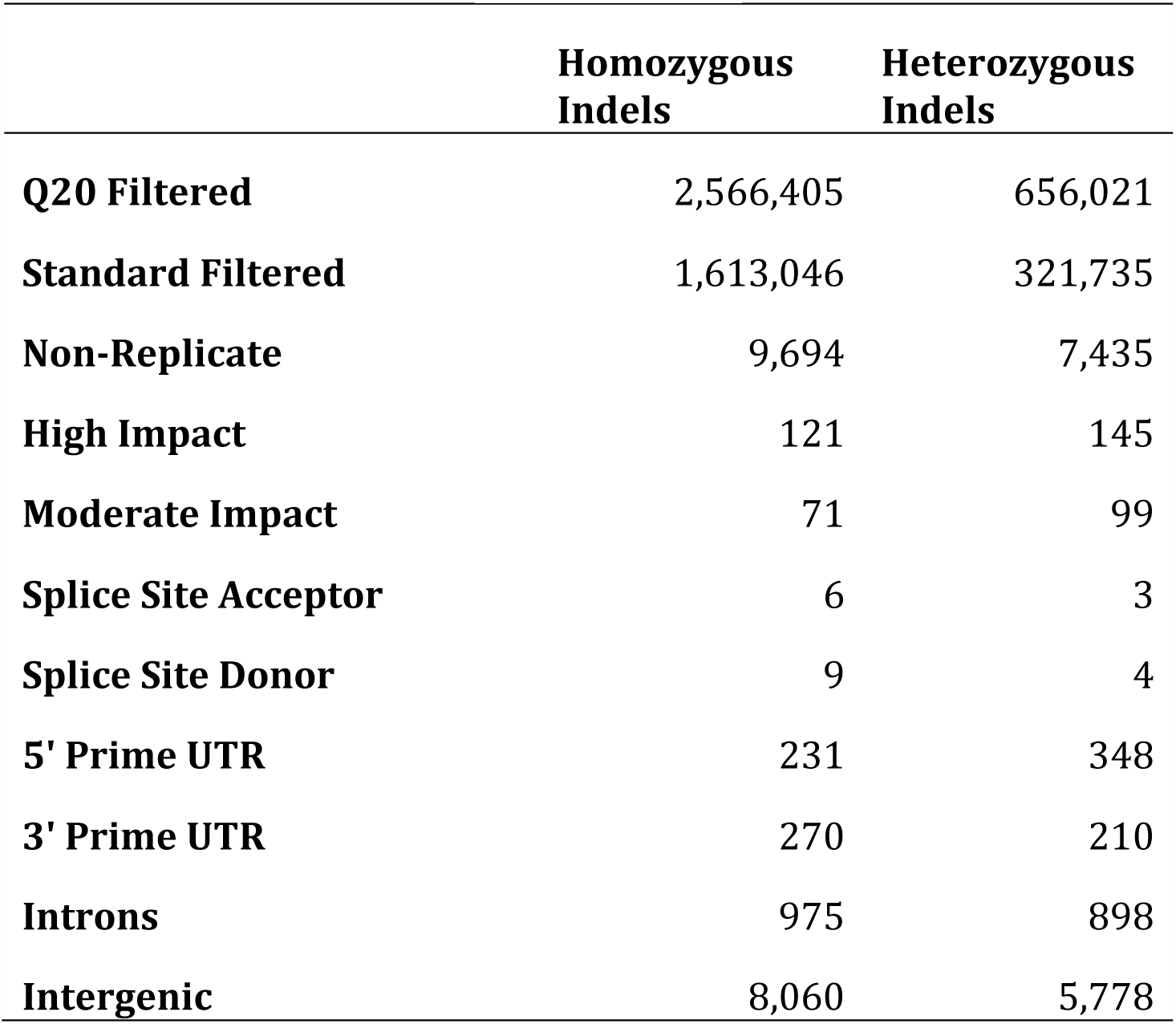
Summary statistics for the detection and annotation of indels in the 570 EMS-mutagenized sorghum BTx623 individuals.

|  | **Homozygous**<br>**Indels** | **Heterozygous**<br>**Indels** |
| --- | --- | --- |
| **Q20 Filtered** | 2,566,405 | 656,021 |
| **Standard Filtered** | 1,613,046 | 321,735 |
| **Non-Replicate** | 9,694 | 7,435 |
| **High Impact** | 121 | 145 |
| **Moderate Impact** | 71 | 99 |
| **Splice Site Acceptor** | 6 | 3 |
| **Splice Site Donor** | 9 | 4 |
| **5' Prime UTR** | 231 | 348 |
| **3' Prime UTR** | 270 | 210 |
| **Introns** | 975 | 898 |
| **Intergenic** | 8,060 | 5,778 |

**Supplementary Table 8.** Summary statistics for the detection and annotation of indels in the 586 EMS-mutagenized sorghum BTx623 individuals.

|  | **Homozygous**<br>**Indels** | **Heterozygous**<br>**Indels** |
| --- | --- | --- |
| **Q20 Filtered** | 2,638,564 | 674,227 |
| **Standard Filtered** | 1,658,457 | 330,621 |
| **Non-Replicates** | 10,074 | 7,686 |
| **High Impact** | 128 | 149 |
| **Moderate Impact** | 75 | 105 |
| **Splice Site Acceptor** | 6 | 3 |
| **Splice Site Donor** | 9 | 4 |
| **5' Prime UTR** | 245 | 358 |
| **3' Prime UTR** | 278 | 213 |
| **Introns** | 1,015 | 932 |
| **Intergenic** | 8,369 | 5,972 |

